# SARS-CoV-2 Nsp2 reprograms host immunity to drive pathogenic inflammation

**DOI:** 10.64898/2026.05.06.723222

**Authors:** Émile Lacasse, Isabelle Dubuc, Joannie Leclerc, Annie Gravel, Leslie Gudimard, Charles Joly Beauparlant, Marion Faure, Patrick Fortin, Marie-Renée Blanchet, Arnaud Droit, Louis Flamand

## Abstract

Despite the end of the COVID-19 pandemic, SARS-CoV-2 continues to circulate endemically, highlighting the need to better understand the viral determinants of pathogenesis. Non-structural protein 2 (Nsp2) has been implicated in host–virus interactions, yet its function remains poorly defined in the context of infection.

Here, we generated a recombinant SARS-CoV-2 lacking Nsp2 (ΔNsp2) to investigate its role in viral replication and disease. While ΔNsp2 replicated comparably to wild-type virus in vitro and in vivo, its deletion resulted in markedly attenuated disease in K18-hACE2 mice.

Wild-type infection induced a strong pro-inflammatory response associated with increased recruitment of monocytes and macrophages, whereas ΔNsp2 infection promoted a more balanced antiviral response characterized by enhanced lymphocyte and NK cell recruitment. This was accompanied by reduced pulmonary and systemic inflammation and distinct transcriptional programs, including downregulation of pathways related to RNA processing and translation.

Mechanistically, CLIP-seq and proximity labeling suggest that Nsp2 interacts with host RNA and components of the translational machinery.

Together, our findings identify Nsp2 as a key virulence factor that drives immunopathology by skewing host immune responses, highlighting its role as a regulator of host–pathogen interactions.

**Graphical Abstract:** 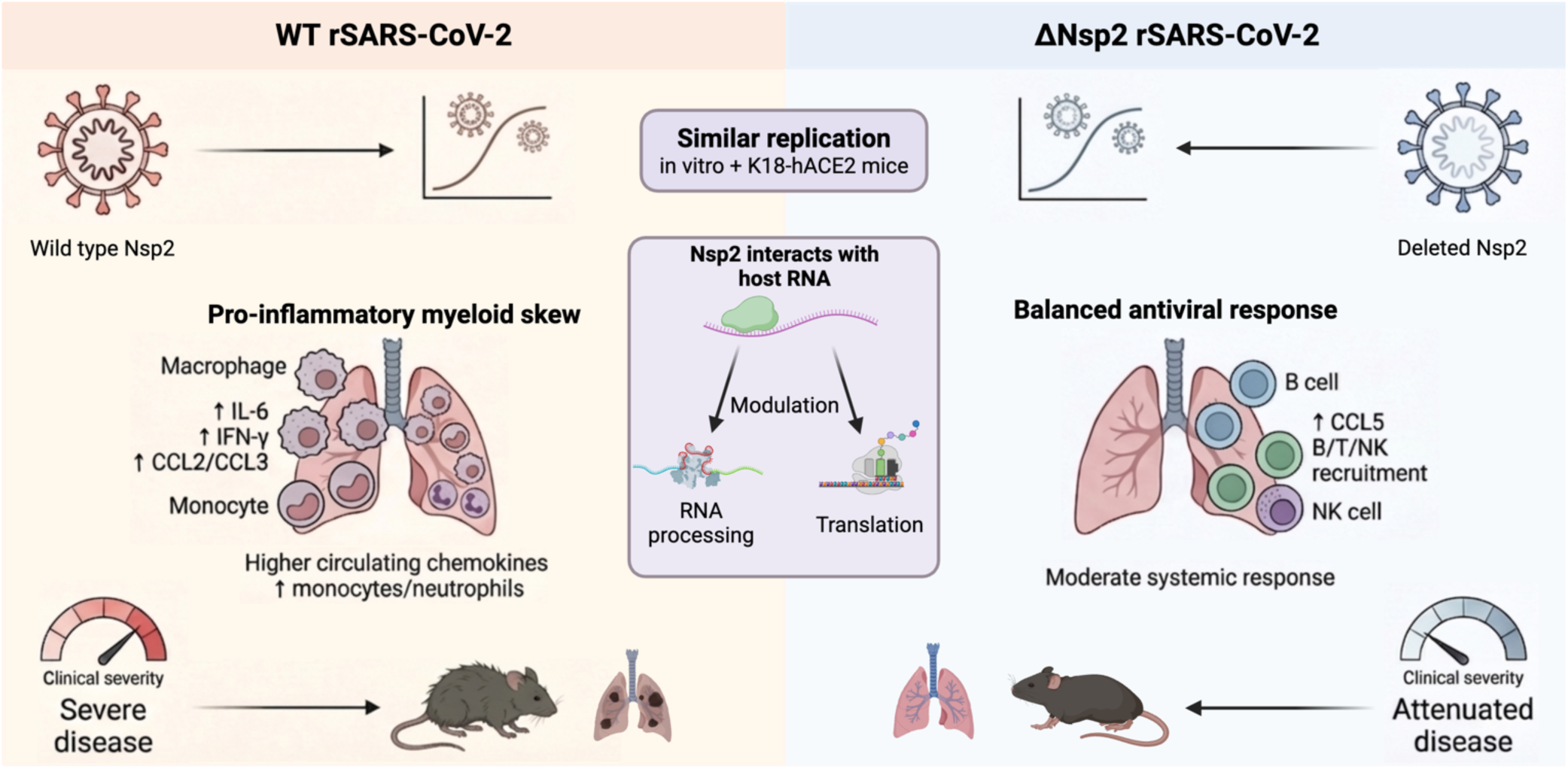

## Introduction

Severe COVID-19 is defined by a dysregulated inflammatory response. Yet, the viral determinants that drive this pathological immune activation remain largely unknown [1–3]. The SARS-CoV-2 genome encodes a large replicase polyprotein that is processed into 16 non-structural proteins (Nsps), several of which actively modulate host innate immune signaling [2, 4].

SARS-CoV-2 infection triggers innate immune sensing through multiple pattern recognition receptors (PRRs), leading to the activation of type I interferon, Nuclear Factor-kappa B (NF-κB), and inflammasome pathways. While these responses are essential for antiviral defense, excessive or prolonged activation drives the production of pro-inflammatory cytokines and chemokines, a hallmark of severe COVID-19 [5–12].

The initiation of these responses results in the production of cytokines, chemokines, and bioactive lipids, leading to profound remodeling of the pulmonary immune cell landscape [13–16]. In patients infected with SARS-CoV-2, the immune response is characterized by elevated circulating levels of proinflammatory cytokines and chemokines, including IL-6, IFN-γ, CCL2 (MCP-1), and CXCL10 (IP-10), indicative of a T helper type 1 (Th1)-polarized response [17–20]. This inflammatory milieu promotes the recruitment of antigen-specific T cells, which contribute to the elimination of virus-infected cells and successful viral clearance [21]. Thus, the same Th1-polarized immune response that promotes viral clearance can, when dysregulated, become a major driver of immunopathology, culminating in acute respiratory syndrome (ARDS) and fatal disease. [21, 22].

Although immune dysfunction is a defining feature of severe SARS-CoV-2 infection, the mechanisms underlying the distinct disease progression observed between common cold coronaviruses and highly pathogenic SARS-related viruses remain poorly understood. Several SARS-CoV-2 proteins have been proposed to play roles in viral pathogenesis [4, 23]. Functional validation of candidate virulence factors remains limited due to the constraints of biosafety level 3 (BSL-3) containment and the technical challenges of coronavirus reverse genetics. Despite the absence of a clearly defined role in viral replication, Nsp2 displays substantial sequence variability among coronaviruses and has been reported to play roles in immune modulation, suggesting a function in host–pathogen interactions rather than genome replication. [24–29]. These observations suggest that Nsp2 may play an underappreciated and critical role in SARS-CoV-2 pathogenesis.

Here, we directly tested the contribution of Nsp2 to SARS-CoV-2 pathogenesis by generating a recombinant virus lacking the entire Nsp2 coding sequence. We show that while Nsp2 is dispensable for viral replication, it is a critical determinant of inflammatory pathology and disease severity *in vivo*. These findings reveal Nsp2 as a previously underappreciated driver of SARS-CoV-2-induced immunopathology and provide a framework for understanding how non-essential viral proteins shape disease severity.

## Materials and methods

### Reverse genetic system

SARS-CoV-2 genomes were assembled into a yeast artificial chromosome (YAC) using transformation-associated recombination (TAR) as described by Thao & al. [30]. Briefly, the YAC vector p0521s gift from Hamilton Smith (Addgene #62862)[31] was amplified with KOD Hot Start Master Mix (MiliporeSigma, Burlington, MA, USA).

The SARS-CoV-2 genome was divided into six overlapping fragments amplified using Platinum™ SuperFi II DNA Polymerase (Thermo Fisher Scientific, Mississauga, ON, Canada) from a Wuhan-like SARS-CoV-2 bacterial artificial chromosome (BAC) kindly provided by Dr. Luis Martinez-Sobrido [32]. Fragments 1 and 6 were subcloned into a pUC19 vector (New England Biolabs [NEB], Whitby, ON, Canada) containing a human cytomegalovirus immediate-early enhancer/promoter, the p0521s insertion site, and a synthetic poly(A) tail followed by a hepatitis delta virus ribozyme sequence, assembled using NEBuilder® HiFi DNA Assembly (NEB).

The fragment encompassing Nsp2 was subcloned into a standard pUC19 vector using NEBuilder® HiFi DNA Assembly. The Nsp2 coding sequence was deleted using the Q5® Site-Directed Mutagenesis Kit (NEB), generating the ΔNsp2 construct.

For TAR assembly, 50 fmol of each SARS-CoV-2 genome fragment and the PCR-linearized vector were co-transformed into Saccharomyces cerevisiae VL6-48N cells [33] using Gietz and Schiestl protocol [34]. Transformed yeast were plated onto synthetic dropout agar lacking arginine, histidine, leucine, tryptophan, and uracil, supplemented with yeast nitrogen base, amino acids, and glucose, and incubated at 30°C for 3 days.

Yeast colonies were lysed in 12 µL of 20 mM NaOH for 10 min at 99°C. Correct assembly of SARS-CoV-2 genome fragments was verified by multiplex PCR using Q5® High-Fidelity DNA Polymerase (NEB). Validated clones were expanded for large-scale YAC purification using the NucleoBond Xtra BAC kit (Macherey-Nagel, Allentown, PA, USA) following a yeast-adapted protocol based on Noskov & al protocol [35].

Briefly, 0.5 L of dropout medium was inoculated with 2 × 10⁸ yeast cells from an 8h preculture and incubated at 30°C with shaking (250 rpm) for 16–18 h until an OD₆₀₀ of ∼16 was reached. Yeast cells were pelleted (5,500 × g, 15 min, 4°C), frozen, thawed, and resuspended in SPE buffer (1 M sorbitol [MiliporeSigma]; 10 mM sodium phosphate [MP Biomedicals, Solon, OH, USA]; 10 mM EDTA [Thermo Fisher Scientific]). Cell walls were digested for 1 h at 37°C using zymolyase 100T (United States Biological, Salem, MA, USA) and β-mercaptoethanol (MiliporeSigma).

Following lysis, DNA purification proceeded according to the NucleoBond Xtra BAC protocol. Lysates were clarified by centrifugation (20,000 × g, 30 min, 4°C) and filtered prior to column purification. Purified YAC DNA was treated with RecBCD exonuclease (5 U/µg DNA) (NEB) in the presence of ATP for 16 h at 37°C, followed by phenol–chloroform–isoamyl alcohol extraction and ethanol precipitation.

YAC and plasmid constructs were sequenced using libraries prepared with the Rapid Barcoding Kit 24 v14 (Oxford Nanopore Technologies, Lexington, MA, USA) and run on Flongle flow cells (Oxford Nanopore Technologies) using a MinION Mk1B (Oxford Nanopore Technologies). Sequencing data were analyzed using the wf-clone-validation pipeline via EPI2ME (Oxford Nanopore Technologies).

### Cell culture and virus rescue

Vero, A549, and A549-hACE2 cells were cultured as described previously [24]. Recombinant SARS-CoV-2 (rSARS-CoV-2) strains were rescued by transfection of purified SARS-CoV-2 constructs into Vero cells stably expressing the SARS-CoV-2 nucleocapsid (N) protein using TransIT-X2® (Chromatographic Specialties Inc., Brockville, ON, Canada), according to the manufacturer’s instructions. A549-hACE2 cells expressing GFP or SARS-CoV-2 Nsp2 (Myc-tagged in at the N-terminus) on the control of a Tet-inducible promoter were generated previously [24].

Supernatants containing progenitor virus were harvested between 3 and 7 days post-transfection based on cytopathic effect. Virus stocks used for mouse experiments were derived from passage 1. Infectious titers were 3.08 × 10⁵ TCID50/mL for the Wuhan-like wild-type (WT) recombinant virus and 5.94 × 10⁵ TCID50/mL for ΔNsp2. For in vitro experiments, the same ΔNsp2 preparation and a WT passage 2 stock (6.81 × 10⁶ TCID50/mL) were used. Recombinant SARS-CoV-2 (rSARS-CoV-2) strains were propagated and titrated on Vero cells as described before [24].

Wuhan-like non-recombinant SARS-CoV2 was obtained from the Laboratoire de Santé Publique du Québec and propagated as described [36, 37]. Infectious titer of the virus sock used was 5.24 × 10⁶ TCID50/mL.

All experiments involving infectious SARS-CoV-2 were conducted under biosafety level 3 conditions. Virus stocks were sequence-validated by RT-PCR using the xGen™ SARS-CoV-2 Midnight Amplicon Panel v2 (Integrated DNA Technologies, Coralville, IA, USA) and analyzed with the wf-artic pipeline via EPI2ME.

### Protein expression

Vero cells were seeded in 6-wells plates at a density of 2.9 × 10^6^ cells per well 24h prior to infection. Cells were infected with recombinant SARS-CoV-2 (rSARS-CoV-2) at a dose of 9.06 × 10^4^ TCID₅₀ for 1 h at 37°C in a humidified incubator with 5% CO₂. After adsorption, cells were washed twice with phosphate-buffered saline (PBS; Corning Life Sciences, Tewksbury, MA, USA) and incubated in complete culture medium.

A549-*hACE2* GFP and SARS-CoV-2 Nsp2 cell lines were seeded in 12-wells plates at a density of 8 × 10^4^ cells per well 24h prior treatment. Protein expression was induced with 0.5µg/mL doxycycline (Takara Bio USA, San Jose, CA, USA).

At 48h post-infection or post-induction, cells were lysed in radioimmunoprecipitation assay (RIPA) buffer (30 mM Tris-HCl, pH 7.5 [Thermo Fisher Scientific];150 mM NaCl [Thermo Fisher Scientific]; 1% Triton X-100 [Thermo Fisher Scientific]; 10% glycerol [Thermo Fisher Scientific]) supplemented with Halt™ Protease Inhibitor Cocktail (Thermo Fisher Scientific) and incubated for 1 h at room temperature with gentle agitation. Lysates were clarified by centrifugation at 14,000 × g for 10 min at 4°C.

Protein concentrations were determined using the Pierce™ BCA Protein Assay Kit (Thermo Fisher Scientific). Equal amounts of protein were resolved on TGX™ Stain-Free gels (Bio-Rad Laboratories [Canada] Ltd., Mississauga, ON, Canada) and transferred onto low-fluorescence PVDF membranes using the Trans-Blot Turbo system (Bio-Rad Laboratories).

Membranes were blocked for 1 h at room temperature in Tris-buffered saline containing 0.1% Tween-20 (TBST) and 5% nonfat dry milk, then incubated overnight at 4°C with the following diluted primary antibodies.

Then, membranes were incubated for 1 h at room temperature with fluorochrome or HRP conjugated secondary antibody. Immunoblots were imaged using a ChemiDoc MP Imaging System (Bio-Rad Laboratories) using fluorescence or chemiluminescent with Clarity Western ECL Substrate (Bio-Rad Laboratories). Total protein staining using Stain-Free Imaging Technology® (Bio-Rad Laboratories) was used as a loading control. All immunoblot experiments were performed at least twice, and one representative blot is shown. Antibody specifications were listed in Supplementary Table 1.

### Viral growth kinetics

A549-*hACE2* cells (8 × 10⁴ cells per well) and Vero cells (2 × 10⁵ cells per well) were seeded in 12-wells plates 24h prior to infection. Differentiated A549 cells were generated by culturing 3.5 × 10⁵ cells on poly-L-lysine–coated 12-mm Transwell inserts (0.4 µm pore size; Corning) under air–liquid interface (ALI) conditions using PneumaCult™-ALI medium (STEMCELL Technologies Canada, Vancouver, BC, Canada) for three weeks, as previously described [38].

Cells were infected with rSARS-CoV-2 at the following doses: 1.2 × 10⁴ TCID50 for A549-*hACE2* cells, 3 × 10⁴ TCID50 for Vero cells, and 6 × 10⁴ TCID50 for ALI cultures (apical compartment). Infection procedures were performed as described in the protein expression section.

Supernatants from A549-*hACE2* and Vero cultures were collected at 16, 24, 48, and 72h post-infection for viral titration. For ALI cultures, 0.5 mL of prewarmed medium was added to the apical compartment and incubated for 30 min at 37°C, after which apical washes were collected for viral titration.

### CCL5 and IL6 in vitro induction and quantification

A549-*hACE2* GFP and SARS-CoV-2 Nsp2 cell lines were seeded and treated as described in the protein expression section. Twenty-four-hour prior experiment endpoint, 1µg Poly I:C (Cytiva, Vancouver, BC, Canada) was transfected with TransIT-mRNA (Chromatographic Specialties Inc.), according to the manufacturer’s instructions.

Supernatants were harvest and mediators were quantified in the supernatant using Human CCL5/RANTES or IL-6 DuoSet ELISA kit (Bio-Techne Canada, Toronto, ON, Canada).

### Cross-linking and Immunoprecipitation Sequencing (CLIP-seq)

A549-*hACE2* GFP and SARS-CoV-2 Nsp2 cell lines were seeded in 15cm petri dish (2.98× 10^6^ cells per dish) 24h prior to infection. Cells were infected with 2.98× 10^6^ TCID50 of SARS-CoV-2 Wuhan per plate following procedure described above. [24, 37]. Once infected GFP or Nsp2 expression was induced with 1µg/mL of doxycycline. Forty-eight hours post-induction and infection, cells were harvested with trypsin/EDTA (Wisent Inc, Saint-Jean-Baptiste, QC, Canada) and crosslinked with 1% formaldehyde for 10min then quenched with 0.25M glycine. Cell lysis and immunoprecipitation were performed as described by Niranjanakumari & al. [39] using Dynabeads Protein G (Thermo Fisher Scientific), murine RNase inhibitor (NEB), an anti-myc antibody (clone 9E10). Bound RNA was purified using Direct-zol RNA Microprep Kit (Zymo Research, Irvine, CA, USA). Libraries were prepared with NEBNext Single Cell/Low Input RNA Library Prep Kit for Illumina (NEB) and sequenced with NovaSeq 6000 (Illumina, San Diego, CA, USA) for approximatively 8 million reads per sample. Whole experiment was performed twice, and the four samples were sequenced in the same run. CLIP-seq analyses were performed with nf-core/clipseq (v1.0.0)[40] using Piranha (v1.2.1)[41] and Paraclu(v9)[42] for peak calling. Myc-immunoprecipitation of GFP (not-tagged) cells were used to remove background peaks. Genes with identified peak by both peak-caller were analysed for functional enrichment using Metascape [43].

### Proximity biotinylation

Nsp2 (non-codon optimized with T85I mutation) and GFP control were cloned into pCW57.1 TurboID-HA-Strep-Tag vector kindly provided by Dr Amélie Fradet-Turcotte using gateway cloning technology. A549(hACE2) were transduced and selected using 1µg/mL puromycin (Wisent Inc). Proximity biotinylation experiment were performed following Agbo & al. protocol[44].

For each condition, cells were plated in 8 15cm petri dish (2.98× 10^6^ cells per dish) 24h prior to infection. Cells were infected with 2.98× 10^6^ TCID50 Wuhan-like non-recombinant virus per plate following the procedure described above. Following infection, protein expression was induced with 1µg/mL (GFP) or 5µg/mL (Nsp2) of doxycycline for 40h in medium with biotin-depleted FBS. Biotinylation was induced by adding 50µM of biotin for 10min at 37°C, 5% CO_2_. Cells were lysed in modified RIPA buffer (Tris-HCl 50mM pH 7.4 [Thermo Fisher Scientific]; 0.9% NaCl [Thermo Fisher Scientific]; 1mM EGTA [MilliporeSigma]; 0.5mM EDTA [Thermo Fisher Scientific]; 1mM MgCl_2_ [Thermo Fisher Scientific]; 1% IGEPAL[MP Biomedicals]; 0.1% SDS [Thermo Fisher Scientific], 0.4% Sodium deoxycholate [MilliporeSigma]) supplemented with Halt™ Protease Inhibitor Cocktail (Thermo Fisher Scientific) and virus was inactivated à 4°C for 24h. Biotinylated proteins were pull down using streptavidin sepharose HP (Avantor, Mississauga, ON, Canada). Identification of interactors was performed by the Plateforme Protéomique du Centre de Génomique de Québec. Identification of potential Nsp2 interactors were performed using the CRAPome database [45] and SAINT pipeline [46]. Analyses were performed on three biological replicates produced from independent experiments.

### Mouse experiments

B6.Cg-Tg(K18-*hACE2*)2Prlmn/J mice were bred and housed at the Centre de Recherche du Centre Hospitalier Universitaire de Québec–Université Laval animal facility under specific pathogen–free conditions. Nine-week-old male and female mice were lightly anesthetized with isoflurane and intranasally infected with 25 µL of M-199 medium containing 3 × 10³ TCID50 of rSARS-CoV-2 (WT and ΔNsp2) or with M-199 medium alone for mock-infected controls.

Mice (n=7/group) were monitored daily for body weight, activity, coat condition, and respiratory distress. Each parameter was scored on a scale from 0 to 3. Animals reaching a score of ≥2 in two or more categories were euthanized according to ethical guidelines.

Additional groups of mice (n=5/group) were euthanized on days 3, 5, or 7 post-infections. Blood was collected by cardiac puncture into citrate-treated tubes. 100µL of blood were taken for leucocyte analysis and remanding to isolate platelet-free plasma. Lungs were harvested and processed as follows: the post-caval lobe of the right lung (∼4 mg) was used for RNA extraction; the entire left lung was used for histological analysis; the superior and middle lobes of the right lung (∼50–70 mg) were used for tissue homogenization; and the inferior lobe (∼70 mg) was used for leukocyte isolation.

### RNA extraction, RT-ddPCR and RNA-seq

Total RNA was extracted from lung tissue using 500 µL of QIAzol reagent (QIAGEN) and homogenized with the Bead Ruptor Elite system (Thermo Fisher Scientific). Homogenates were centrifuged at 16,000 × g for 5 min at 4°C, and RNA was purified from the supernatant using the Direct-zol RNA Miniprep Plus Kit (Zymo Research) with on-column DNase I treatment.

Viral RNA quantification was performed by reverse transcription droplet digital PCR (RT-ddPCR) as previously described [47]. RNAseq was performed following ribosomal RNA depletion as detailed elsewhere [47].

### RNAseq and statistical analysis

RNA sequencing (RNA-seq) pre-processing steps were performed on the Narval high-performance computing cluster (École de technologie supérieure, Montréal, QC, Canada) provided by the Digital Research Alliance of Canada. Raw sequencing reads were quality-filtered, trimmed, and deduplicated using fastp (v1.0.1) [48]. Processed reads were pseudo-aligned and quantified using kallisto (v0.51.1) [49].

A hybrid reference transcriptome was constructed by combining mouse transcripts from GENCODE (release M36), human ACE2 transcripts, and SARS-CoV-2 transcripts derived from assembly ASM985889v3 with annotations from Ensembl [50]. RNA-seq data from mock-infected K18-hACE2 mice generated in a previous study were included as controls to assess infection-induced gene expression changes [51].

Downstream statistical analyses and data visualization were performed in R (v4.5.2) using RStudio (v2026.01.1+403). Transcript-level abundance estimates were imported using the tximport package (v1.38.2) [52], and differential gene expression analysis was conducted using DESeq2 (v1.50.2) [53]. Functional enrichment analyses were performed using msigdbr (v25.1.1) [54–56] and clusterProfiler (v4.18.4) [57].

Data visualization was carried out using the following R packages: tidyverse (v2.0.0), ComplexHeatmap (v2.26.1) [58], circlize (v0.4.17) [59], ggrepel (v0.9.7), gtools (v3.9.5), grid (v4.5.2), umap (v0.2.10.0), igraph (v2.2.2)[60], ggraph (v2.2.2)[60], scales (v1.4.0) and colorspace (v2.1-2)[61].

Heatmaps used to visualize lung cytokine modulation and GSEA were generated using ComplexHeatmap (v2.26.1) [58]. Additional figures were produced using GraphPad Prism (v11.0.0).

Statistical analyses for all datasets other than RNA-seq were performed using GraphPad Prism. Depending on the experimental design, comparisons were conducted using two-way ANOVA and uncorrected Fisher’s least significant difference (LSD) test.

### Infectious viral load and cytokines quantification

Superior and middle lung lobes were homogenized in 1.5 mL PBS using the Bead Ruptor Elite system and centrifuged at 3,000 × g for 20 min at 4°C. Supernatants were stored frozen (−80°C) until infectious virus titration.

For cytokine and chemokine measurements, lung homogenates and platelet-free plasma samples were treated with 1% Triton X-100 for 1 h at room temperature to inactivate SARS-CoV-2. Cytokines, chemokines, and interferons were quantified using ProcartaPlex™ Mouse Mix & Match 13-plex panels (Life Technologies Inc., Burlington, ON, Canada) on a Bio-Plex 200 instrument (Bio-Rad Laboratories). Total protein concentrations were measured using the Pierce™ BCA Protein Assay (Thermo Fisher Scientific) and used for normalization

### Histological Analysis

Lung tissues were fixed in formalin, embedded in paraffin, sectioned, deparaffinized, and rehydrated using standard procedures [62]. Lung inflammation was assessed on sections stained using the Carstairs (Electron Microscopy Sciences, Hatfield, PA, USA) method. Slides were digitized using an AxioScan Z1 slide scanner (Carl Zeiss, Oberkochen, Germany) at 20× magnification. Quantitative analysis of lung inflammation was performed as previously described [47].

### Leukocyte isolation and flow cytometry

Leukocyte isolation protocol was performed according to Rothfuchs et al[63–65]. Briefly, inferior lung lobes were minced in RPMI 1640 medium and digested with 150 µg Liberase™ TM (MilliporeSigma) and DNase I (MilliporeSigma) at 37°C for 30 min. Digested tissues were passed through 100-µm CellTrics™ filters centrifuged and 500 × g for 5 min. Cell pellets were resuspended in 33% isotonic Percoll-PBS (MilliporeSigma) and centrifuged at 1,000 × g for 20 min at room temperature with slow acceleration and deceleration. Pellets were subjected to ACK red blood cell lysis, washed, and resuspended for staining.

Blood samples were similarly treated with ACK buffer to remove red blood cells. Cells were stained with 0.5µL Ghost Dye™ Red 710 viability dye (Tonbo Biosciences, San Diego, CA, USA), followed by Fc receptor blocking and antibody staining following manufacturer’s instructions. Samples were fixed in 1% paraformaldehyde and analyzed on a Northern Lights™ flow cytometer (Cytek Biosciences, Fremont, CA, USA). Antibody panels and gating strategies are provided in Supplementary Table 2 and Supplementary Figure 1.

## Results

### The SARS-CoV-2 ΔNsp2 mutant displays comparable replication kinetics to WT SARS-CoV-2 in vitro

A Wuhan-like recombinant SARS-CoV-2 and a mutant virus lacking the Nsp2 coding sequence (ΔNsp2) were generated using the reverse genetics system schematized in Figure 1A. Expression of Nsp1, Nsp2 and Nsp3, which are produced by proteolytic cleavage of the ORF1a/b polyprotein, is shown in Figure 1B. When normalized to nucleocapsid (N) protein expression, comparable levels of Nsp1 and Nsp3 were detected in cells infected with either virus, indicating preserved polyprotein processing. As anticipated, Nsp2 expression was absent in cells infected with the ΔNsp2 mutant.

**Figure 1.**
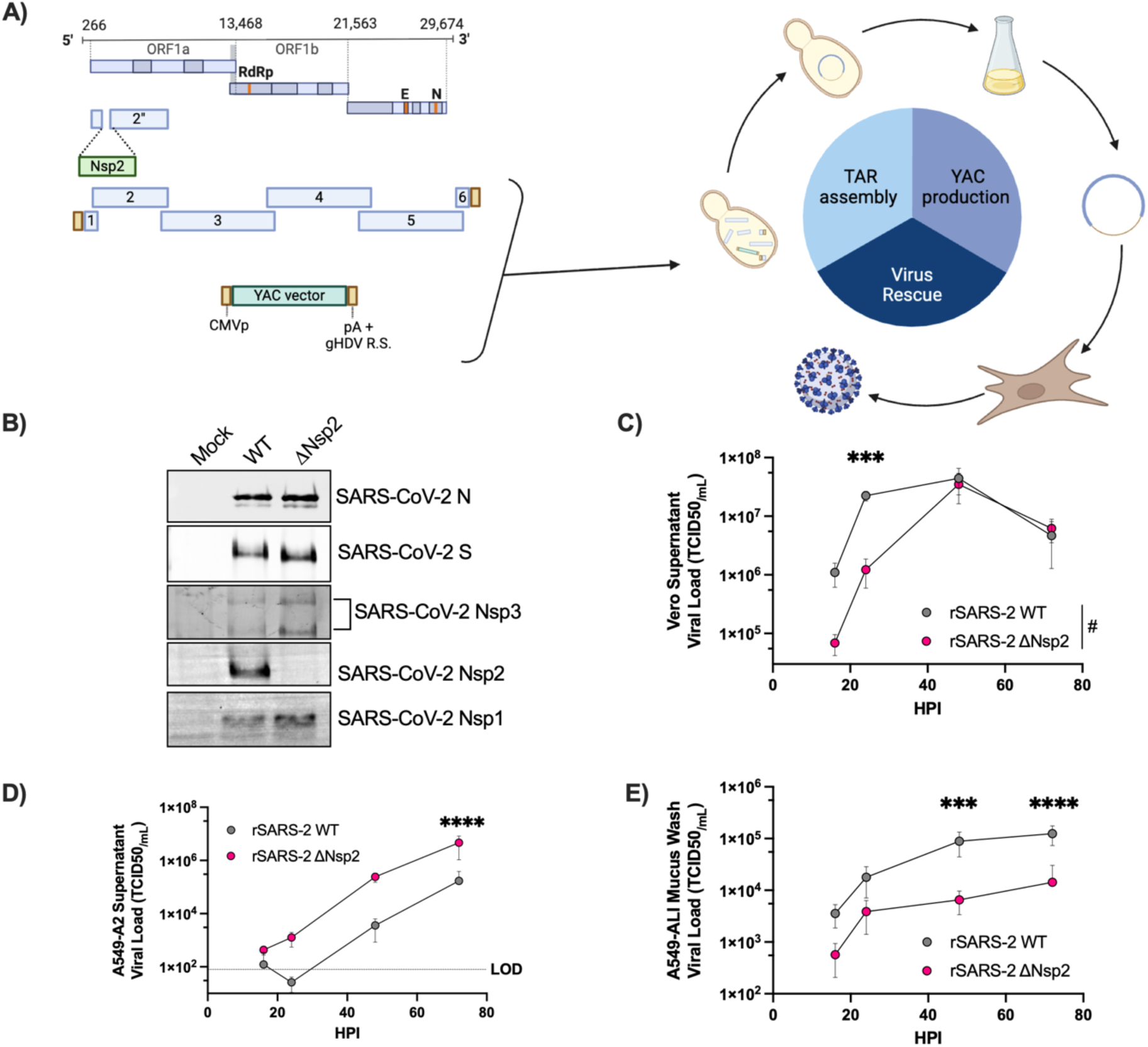
Generation of recombinant SARS-CoV-2 viruses and in vitro growth kinetics. (A) Schematic representation of the SARS-CoV-2 reverse genetics platform used to generate the wild-type (WT) and ΔNsp2 recombinant viruses. The figure was created using BioRender. Overlapping genomic fragments were assembled and recombined into the YAC vector through TAR cloning. YAC DNA was nucleofected into Vero cells to reconstitute the viruses. (B) Viral protein expression following infection of Vero cells with WT or ΔNsp2 recombinant viruses. Cells were infected and harvested as described in the Materials and Methods section. Protein expression was analyzed by Western blot using antibodies against SARS-CoV-2 N (AF488), S(AF647), Nsp3(AF647), Nsp2(AF488) and Nsp1(AF488). Stain-free technology was used as a loading control (see Supplementary Figure 2). The experiment was performed twice, and a representative result is shown. (C–E) Growth kinetics of recombinant viruses in Vero (C), A549-*hACE2* (D), and A549-ALI (E) cells. Viral shedding was quantified in cell culture supernatants (C and D) or mucus washes (E). Viral titers are expressed as TCID50/mL (mean ± SD; n = 3–6 per group). The dashed line indicates the limit of detection (LOD). The x-axis represents hours post-infection (hpi). Global growth kinetics were compared using two-way ANOVA. ^#^P < 0.033. Individual time points were analyzed using uncorrected Fisher’s LSD test. ***P < 0.0002; ****P < 0.000

Viral replication was next assessed using three in vitro infection models: African green monkey kidney cells (Vero), human alveolar type II–like carcinoma cells overexpressing human *ACE2* (A549-*hACE2*), and differentiated A549 cells cultured at the air–liquid interface (A549-ALI). As shown in Figures 1C–E, the wild-type and ΔNsp2 viruses exhibited largely comparable growth kinetics across all models. While peak viral titers varied depending on the cellular system, no consistent replication defects were observed for the ΔNsp2 mutant. Wild-type recombinant virus reached higher titers in Vero and A549-ALI cultures, the ΔNsp2 mutant achieved higher titers in A549-*hACE2* cells.

### The ΔNsp2 mutant induces an attenuated disease in K18-hACE mice

K18-*hACE2* transgenic mice were infected intranasally with equal quantity of infectious wild-type or ΔNsp2 recombinant SARS-CoV-2, as outlined in Figure 2A. Predefined groups of animals were euthanized at specific time points for tissue collection, while the remaining mice were monitored daily for clinical signs of disease. Strikingly, the majority of mice infected with the ΔNsp2 mutant recovered from infection and survived until the experimental endpoint (Figure 2B). In contrast, infection with the wild-type virus resulted in a markedly worse outcome, with only approximately 14% of mice surviving beyond day 8 post-infection (DPI).

**Figure 2.**
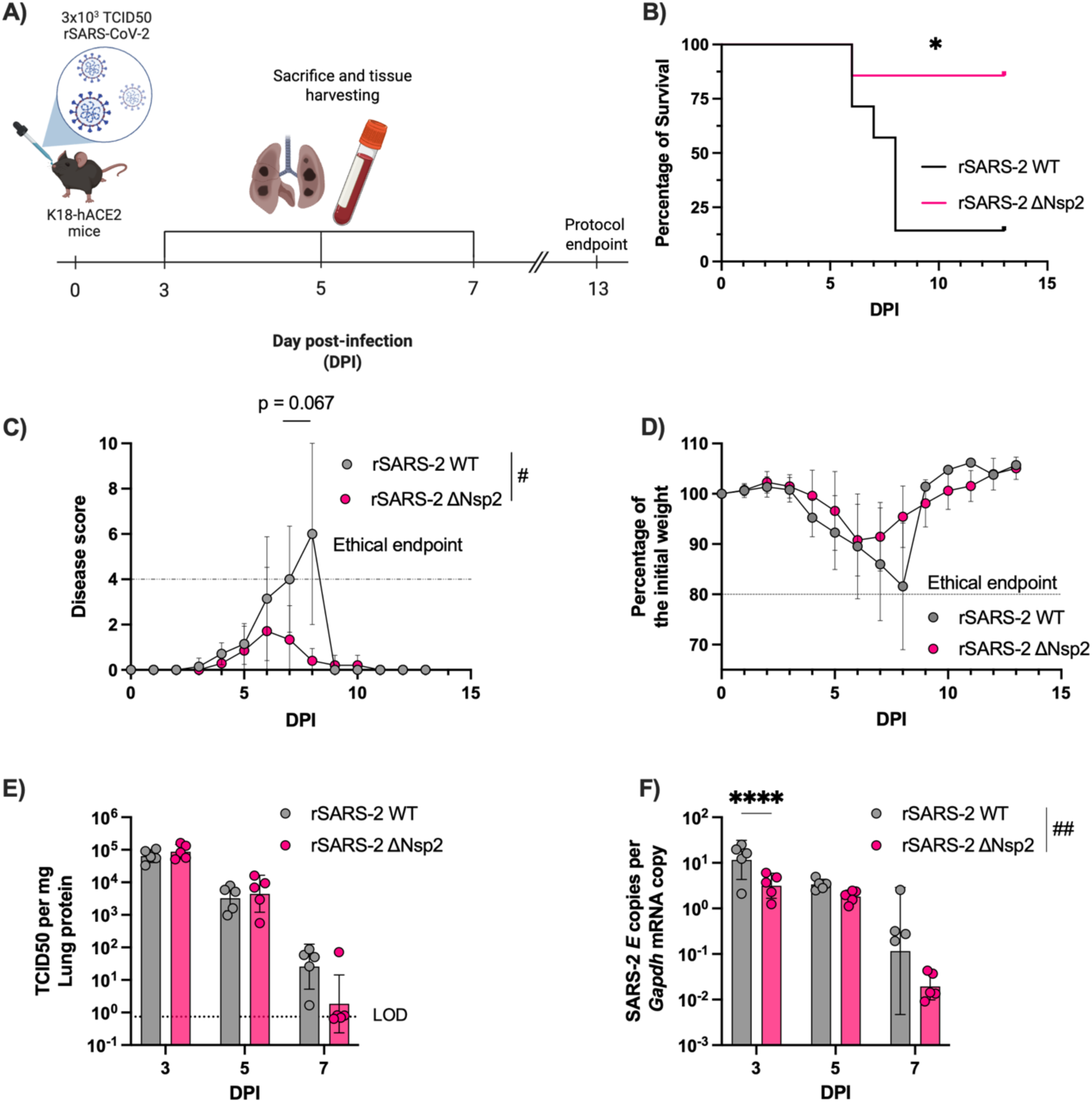
Attenuated disease severity in K18-*hACE2* mice infected with ΔNsp2 recombinant SARS-CoV-2. (A) Schematic overview of the mouse infection protocol. This figure was created using BioRender. (B) Survival curves of K18-*hACE2* mice following infection with wild-type or ΔNsp2 recombinant SARS-CoV-2. Survival is expressed as the percentage of mice not reaching the ethical endpoint at each day post-infection (DPI) (n = 7 per group). Statistical analysis was performed using the Mantel–Cox (log-rank) test. *P < 0.033. (C) Clinical disease scores following infection. Disease scores represent the cumulative score of individual parameters including body weight, activity, coat condition, and respiratory symptoms (each scored from 0 to 3). A total score of 4 corresponded to the ethical endpoint, as euthanasia was required when two or more parameters reached a score ≥2 (mean ± SD; n = 7 per group). (D) Body weight changes following infection, expressed as the percentage of initial body weight at each DPI (mean ± SD; n = 7 per group). For panels C and D, differences over the entire infection course were analyzed using mixed-effects models. ^#^P < 0.033. Individual time points were compared using uncorrected Fisher’s LSD tests. (E) Infectious viral loads in lung tissues. Recombinant SARS-CoV-2 titers were determined in lung homogenates using the 50% tissue culture infectious dose (TCID₅₀) assay and normalized to total protein content (mean ± SD; n = 5 per group). (F) SARS-CoV-2 E gene RNA levels in lung tissues, quantified by RT-ddPCR and normalized to mouse Gapdh mRNA copy number (mean ± SD; n = 5 per group). For panels E and F, data across the infection course were analyzed using two-way ANOVA. ^##^P < 0.0021. Groups at individual time points were compared using uncorrected Fisher’s LSD tests. ****P < 0.0001.

Consistent with these survival data, infection with the ΔNsp2 mutant was associated with a significantly reduced disease severity, as reflected by lower clinical scores peaking between days 6 and 9 post-infection (Figure 2C). Mice infected with the ΔNsp2 virus also exhibited a trend toward reduced body weight loss compared to wild-type–infected animals, although this difference did not reach statistical significance (Figure 2D).

To determine whether disease attenuation was associated with impaired viral replication in vivo, infectious viral loads were measured in lung tissues. While no statistically significant differences in lung infectious titers were observed between the two groups, a trend toward more efficient viral clearance was detected in ΔNsp2-infected mice at 7 DPI (Figure 2E). In parallel, quantification of viral RNA revealed an overall reduction in SARS-CoV-2 E gene expression in the lungs of ΔNsp2-infected mice, which was most pronounced at 3 DPI (Figure 2F).

### Wild-type SARS-CoV-2 infection elicits a stronger inflammatory response in the lung

To characterize the pulmonary inflammatory response induced by wild-type and ΔNsp2 recombinant SARS-CoV-2, a panel of 13 immune mediators, including cytokines, chemokines, and interferons, were quantified in lung homogenates (Figure 3A). At early time points, levels of the monocyte-recruiting chemokine CCL2 (MCP-1) were significantly higher in lungs from wild-type–infected mice at 3 days post-infection (DPI). At 5 DPI, wild-type infection was associated with increased production of CXCL9 (MIG) and IFNγ, indicative of a heightened type 1 inflammatory response.

**Figure 3.**
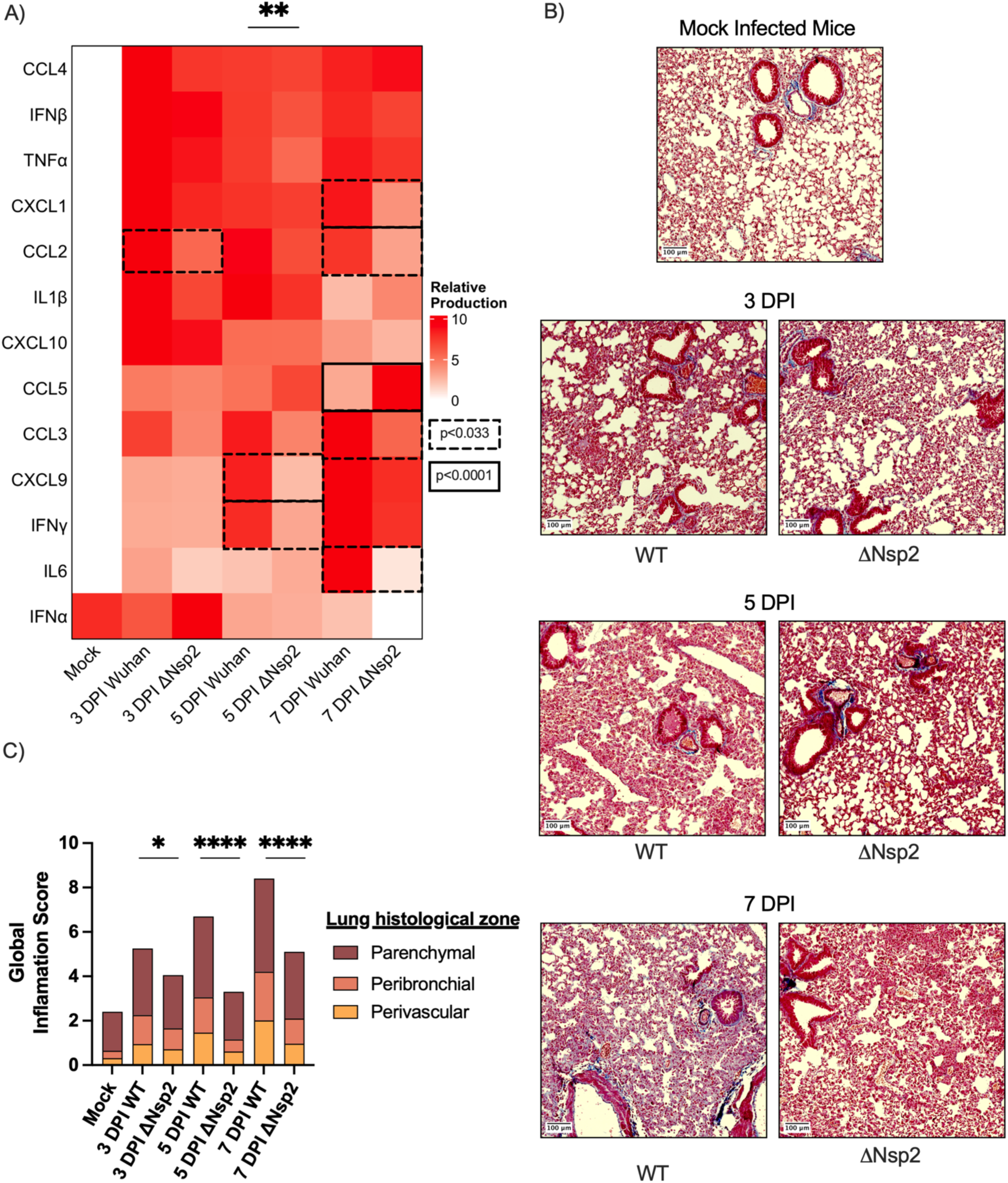
Enhanced pulmonary inflammation following wild-type rSARS-CoV-2 infection. (A) Cytokine, chemokine, and interferon profiles in lung tissues following mock or rSARS-CoV-2 infection. Mediator concentrations were normalized to total protein content, and values were scaled from 0 to 10 for each mediator to facilitate visualization in the heatmap (mean; n = 5 per group). Differences between groups at the same time point for the mediator profile were assessed using two-way ANOVA (**P < 0.0021), followed by Fisher’s LSD test for individual mediators. (B) Carstairs staining of lung sections from mock-, wild-type–, or ΔNsp2-infected K18-*hACE2* mice. Whole lung sections were scanned, and representative images from each group are shown. (C) Histological assessment of lung inflammation in perivascular, peribronchial, and parenchymal regions. Whole lung sections were independently scored for inflammation (scale 0–5) in each anatomical compartment (mean; n = 4–5 per group). Global inflammation across time points was analyzed using Fisher’s LSD test. (C) *P < 0.033; ****P < 0.0001

At later stages of infection (7 DPI), several pro-inflammatory mediators, including CXCL1 (KC), CCL2, CCL3 (MIP-1α), and IL-6, were detected at significantly higher levels in lungs from wild-type–infected mice compared to ΔNsp2-infected animals. In contrast, the T cell–associated chemokine CCL5 (RANTES) was more abundant in lungs from ΔNsp2-infected mice at 7 DPI. Reduction of CCL5 production in the presence of Nsp2 was partially recapitulated in A549-*hACE2* stimulated with Poly I:C (Supplementary Figure 4B). Under the same conditions, Nsp2 did not influence IL6 production (Supplementary Figure 4C).

These findings were correlated with histopathological changes. Lung sections from wild-type–infected mice exhibited more pronounced inflammatory infiltrates in perivascular, peribronchial, and parenchymal regions compared to lungs from ΔNsp2-infected animals (Figures 3B and 3C), consistent with exacerbated lung inflammation following wild-type rSARS-CoV-2 infection.

### Differential modulation of lung leukocyte populations following infection with wild-type or ΔNsp2 rSARS-CoV-2

Leukocytes were isolated from harvested lung lobes and analyzed by flow cytometry using the gating strategy described in Supplementary Figure 1. Total leukocyte recruitment in the lungs was significantly increased at 7 days post-infection (DPI) for both recombinant viruses, with no statistically significant difference between groups (Figure 4A). However, comprehensive immunophenotyping revealed marked differences in lung leukocyte sub-populations between wild-type– and ΔNsp2-infected mice (Figure 4B). A distinct global leukocyte profile was already evident at 5 DPI, indicating divergent immune responses elicited by the two viruses.

**Figure 4.**
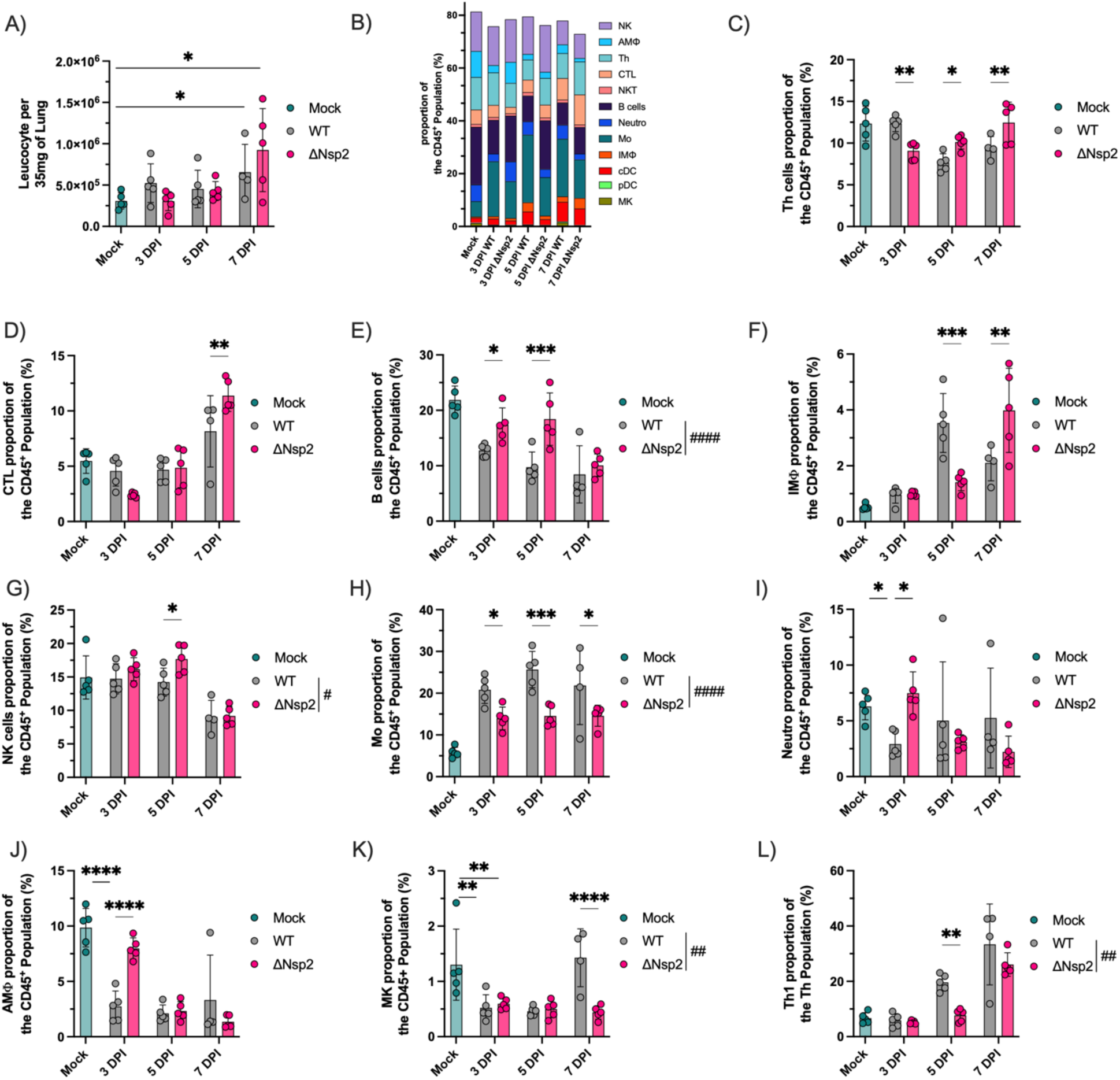
Differential modulation of lung leukocyte populations following infection with wild-type or ΔNsp2 mutant rSARS-CoV-2. (A) Total lung leukocyte counts in mock- and rSARS-CoV-2–infected mice. Inferior lung lobes were dissociated, and leukocytes were isolated and stained for flow cytometry as described in the Materials and Methods section. Data are expressed as absolute leukocyte counts normalized to the weight of the corresponding lung tissue (half of the inferior lobe, ∼35 mg) (mean ± SD; n = 4–5 per group). (B) Global leukocyte composition in the lung. Each colored bar represents the proportion of each leukocyte subset within total CD45⁺ cells (mean; n = 4–5 per group). (C–L) Proportions of individual leukocyte populations among total CD45⁺ lung cells: T helper (Th) cells (C), cytotoxic T lymphocytes (CTLs) (D), B cells (E), interstitial macrophages (IMΦ) (F), natural killer (NK) cells (G), monocytes (Mo) (H), neutrophils (Neutro) (I), alveolar macrophages (AMΦ) (J), megakaryocytes (MK) (K), and T helper type 1 (Th1) cells (L). Data are presented as percentages of CD45⁺ cells (mean ± SD; n = 4–5 per group). Comparisons across the infection course were performed using two-way ANOVA. ^#^P < 0.033; ^##^P < 0.0021; ^####^P < 0.0001. Group comparisons at individual time points were assessed using uncorrected Fisher’s LSD test. *P < 0.033; **P < 0.0021; ***P < 0.0002; ****P < 0.0001.

ΔNsp2 infection was associated with increased proportions of adaptive immune cells. Specifically, higher frequencies of T helper (Th) cells and B cells were observed at 5 and 7 DPI, cytotoxic T lymphocytes (CTLs) at 7 and B cells throughout the course of infection compared to wild-type infection (Figures 4C–4E).

In contrast, innate immune cell populations exhibited distinct kinetics depending on the viral strain. Interstitial macrophage (IMΦ) accumulation peaked earlier following wild-type infection (5 DPI), whereas IMΦ recruitment was delayed and peaked at 7 DPI in ΔNsp2-infected lungs (Figure 4F). Natural killer (NK) cells were more abundant in ΔNsp2-infected lungs, with the most pronounced difference observed at 5 DPI (Figure 4G). Conversely, monocyte (Mo) proportions were consistently higher across all time points in wild-type–infected lungs (Figure 4H).

Neutrophil and alveolar macrophage (AMΦ) populations were transiently reduced at 3 DPI following wild-type infection but not after ΔNsp2 infection (Figures 4I and 4J). Lung megakaryocyte (MK) abundance was also reduced following infection; however, recovery at 7 DPI was observed only in wild-type–infected mice (Figure 4K).

Finally, analysis of T helper cell differentiation revealed a higher proportion of Th1 cells in lungs from wild-type–infected mice compared to ΔNsp2-infected animals (Figure 4L), consistent with the enhanced type 1 inflammatory environment induced by the wild-type virus.

### Shift in antigen-presenting cell populations in the lung following wild-type or ΔNsp2 rSARS-CoV-2 infection

Analysis of antigen-presenting cell populations revealed marked differences between wild-type– and ΔNsp2-infected lungs. As shown in Figures 5A and 5B, classical dendritic cells (cDCs) and plasmacytoid dendritic cells (pDCs) accounted for a significantly greater proportion of total lung leukocytes following infection with wild-type rSARS-CoV-2 compared to ΔNsp2 mutant infection.

**Figure 5.**
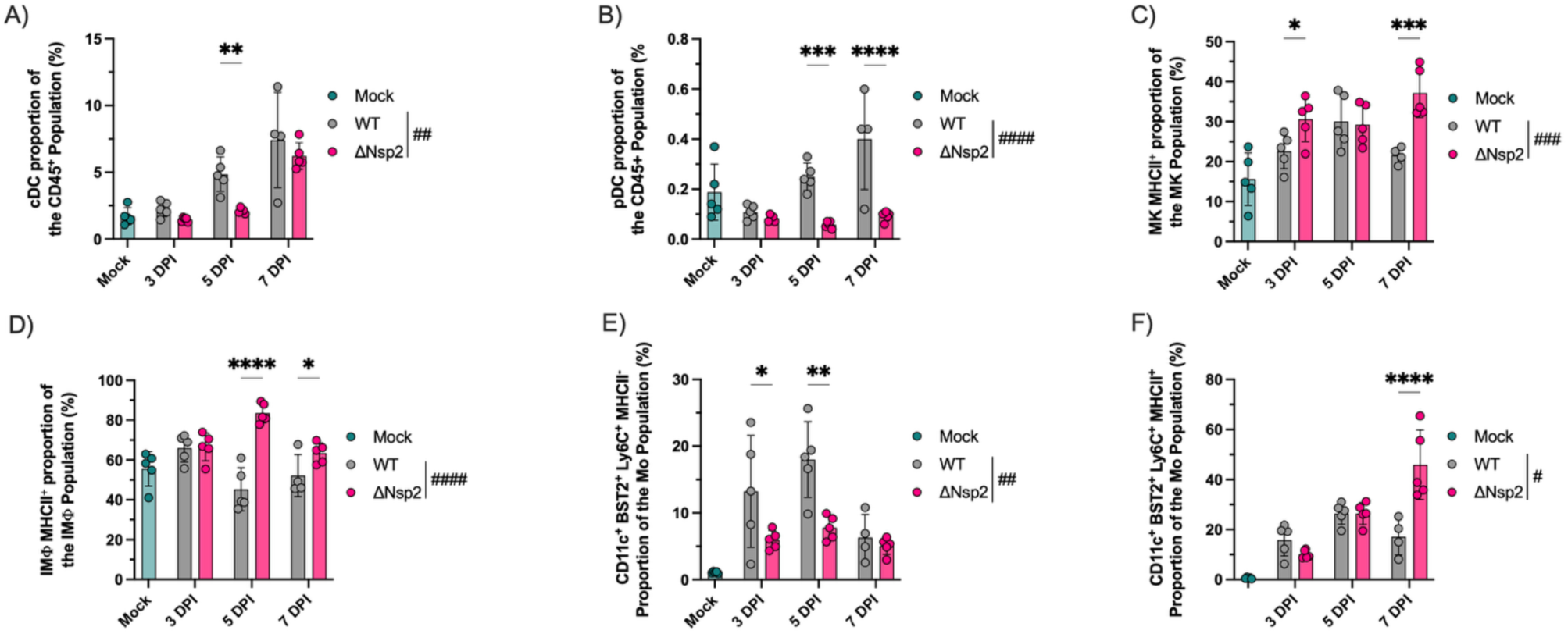
Modulation of antigen-presenting cell populations in the lung following infection with wild-type or ΔNsp2 mutant rSARS-CoV-2. (A–B) Abundance of classical dendritic cells (cDCs) (A) and plasmacytoid dendritic cells (pDCs) (B) expressed as a percentage of total CD45⁺ lung leukocytes (mean ± SD; n = 4–5 per group). (C) Proportion of MHC class II (MHCII)–positive cells within the lung megakaryocyte (MK) population (mean ± SD; n = 4–5 per group). (D) Modulation of MHCII expression on interstitial macrophages (IMΦ), expressed as the percentage of MHCII⁺ cells within the total IMΦ population (mean ± SD; n = 4–5 per group). (E–F) Proportions of monocytes (Mo) co-expressing CD11c, BST2, and Ly6C that were MHCII-positive (E) or MHCII-negative (F), expressed as percentages of the total monocyte population (mean ± SD; n = 4–5 per group). Comparisons across the infection course were performed using two-way ANOVA. ^#^P < 0.033; ^##^P < 0.0021; ^###^P < 0.0002; ^####^P < 0.0001. Group comparisons at individual time points were assessed using uncorrected Fisher’s LSD test. *P < 0.033; **P < 0.0021; ***P < 0.0002; ****P < 0.0001.

In contrast, ΔNsp2 infection was associated with enhanced expression of major histocompatibility complex class II (MHCII) on non-canonical antigen-presenting cell populations. Specifically, a higher proportion of lung megakaryocytes (MKs) and interstitial macrophages (IMΦs) expressed MHCII in ΔNsp2-infected mice relative to wild-type–infected animals (Figures 5C and 5D).

Similarly, monocyte subsets co-expressing CD11c, bone marrow stromal cell antigen 2 (BST2), and lymphocyte antigen 6C (Ly6C) displayed increased MHCII expression in response to ΔNsp2 infection compared to wild-type infection (Figures 5E and 5F). Together, these data indicate that deletion of Nsp2 profoundly alters the landscape of antigen presentation in the lung, shifting MHCII expression away from classical dendritic cells toward alternative myeloid and stromal-associated populations.

### ΔNsp2 infection elicits a robust antiviral response in peripheral blood, whereas wild-type rSARS-CoV-2 promotes a pro-inflammatory systemic profile

As in the lung, chemokines, interferons, and leukocyte populations were quantified in blood or plasma from mock-, wild-type–, and ΔNsp2-infected mice. A more pronounced systemic chemokine and interferon response was detected in mice infected with wild-type rSARS-CoV-2 compared to those infected with the ΔNsp2 mutant (Figure 6A). Notably, the inflammatory burst in plasma occurred earlier during wild-type infection, with significant differences between mock and wild-type groups detected at 5 DPI (P = 0.0001), whereas a delayed response was observed in ΔNsp2-infected mice, reaching significance only at 7 DPI (P = 0.018).

**Figure 6.**
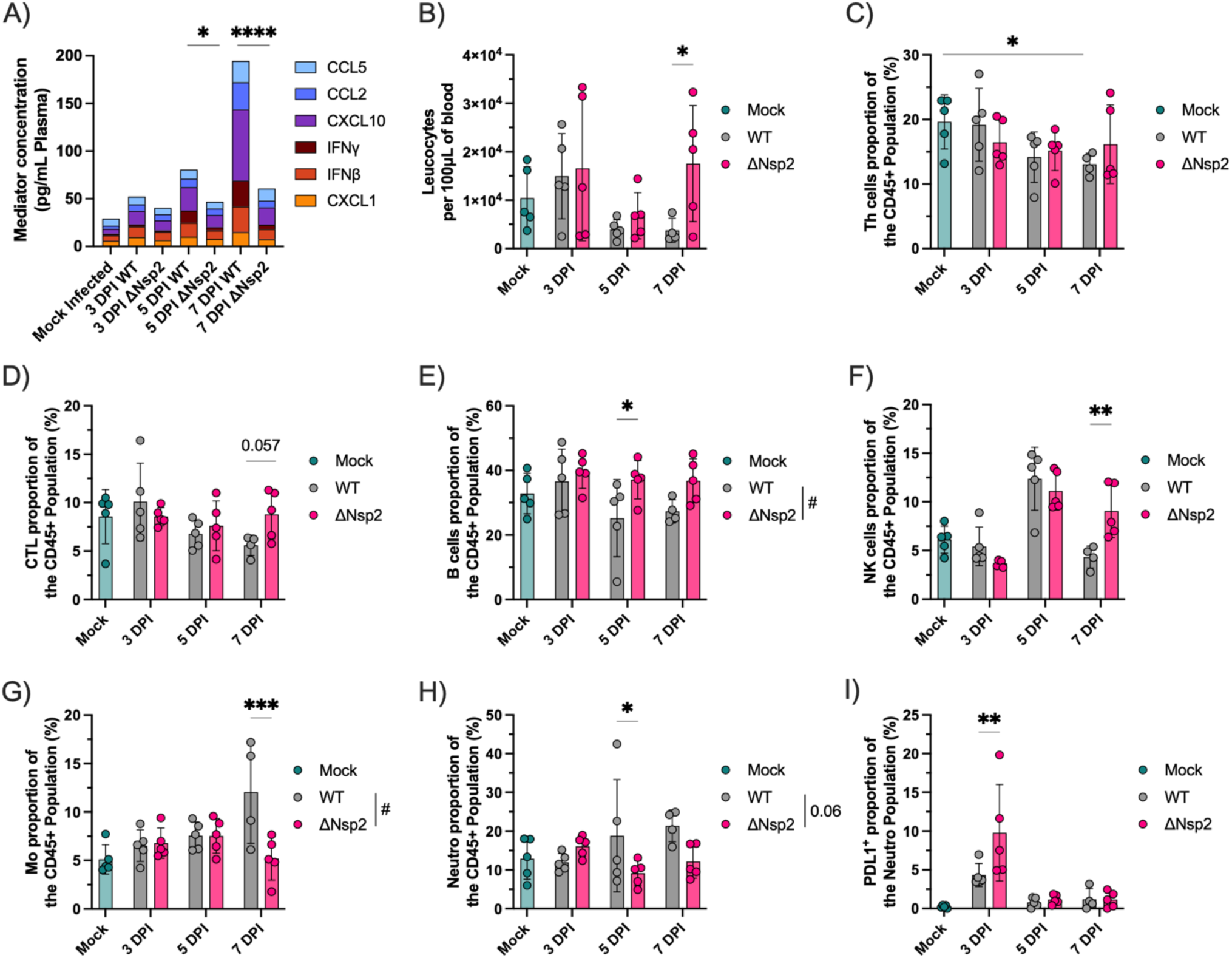
Systemic immune responses following mock, wild-type, or ΔNsp2 rSARS-CoV-2 infection in K18-*hACE2* mice. (A) Systemic inflammatory response in plasma from mock-, wild-type–, or ΔNsp2-infected mice. Chemokines and interferons were quantified in Triton X-100–inactivated, platelet-depleted plasma and expressed as pg/mL (mean; n = 4–5 per group). Mean mediator levels were compared between groups at the same time point using uncorrected Fisher’s LSD test. (B) Quantification of circulating leukocytes. Citrate-treated whole blood was subjected to red blood cell lysis, and leukocytes were stained and analyzed by flow cytometry. Results are expressed as CD45⁺ cells per 100 µL of whole blood. (C–I) Abundance of circulating T helper (Th) cells (C), cytotoxic T lymphocytes (CTLs) (D), B cells (E), natural killer (NK) cells (F), monocytes (Mo) (G), neutrophils (Neutro) (H), and PD-L1–expressing neutrophils (I). Data are presented as percentages of CD45⁺ cells (mean ± SD; n = 4–5 per group). For all panels, comparisons across the infection course were performed using two-way ANOVA. ^#^P < 0.033. Group comparisons at individual time points were assessed using uncorrected Fisher’s LSD test. *P < 0.033; **P < 0.0021; ***P < 0.0002; ****P < 0.0001.

Additional cytokines, chemokines, and interferons measured in lung homogenates were also assessed in plasma; however, their concentrations were either close to the limit of detection or did not significantly differ between infection groups.

Total circulating leukocyte counts did not differ markedly between mock- and virus-infected mice throughout the course of infection (Figure 6B). However, at 7 DPI, higher leukocyte counts were observed in mice infected with the ΔNsp2 mutant compared to wild-type–infected animals.

Wild-type infection was associated with a tendency toward peripheral T cell depletion. While reductions were observed for both T helper (Th) cells and cytotoxic T lymphocytes (CTLs), only the decrease in Th cells at 7 DPI reached statistical significance (Figures 6C and 6D). In contrast, ΔNsp2 infection was associated with increased B cell abundance in the blood, most prominently at 5 DPI (Figure 6E).

Natural killer (NK) cell proportions increased at 5 DPI following infection with both viruses; however, this increase persisted to 7 DPI only in ΔNsp2-infected mice (Figure 6F). Conversely, wild-type infection was characterized by increased proportions of circulating monocytes at 7 DPI and neutrophils at 5 DPI (Figures 6G and 6H).

Within the neutrophil compartment, the proportion of cells expressing the inhibitory receptor programmed death-ligand 1 (PD-L1) was higher at 3 DPI in mice infected with the ΔNsp2 mutant compared to wild-type–infected animals (Figure 6I), suggesting an early regulatory phenotype associated with attenuated disease.

### Nsp2 deletion alters early lung transcriptional responses to SARS-CoV-2 infection

RNA extracted from infected mouse lungs was sequenced and transcripts relative abundance was quantified. Dimensionality reduction of variance-stabilized counts revealed a clear separation between wild-type and ΔNsp2-infected mice at 3 days post-infection (DPI), indicating distinct early transcriptional responses (Figure 7A). A similar trend was observed at 5 DPI, although increased variability within the wild-type group led to the emergence of two subclusters.

**Figure 7.**
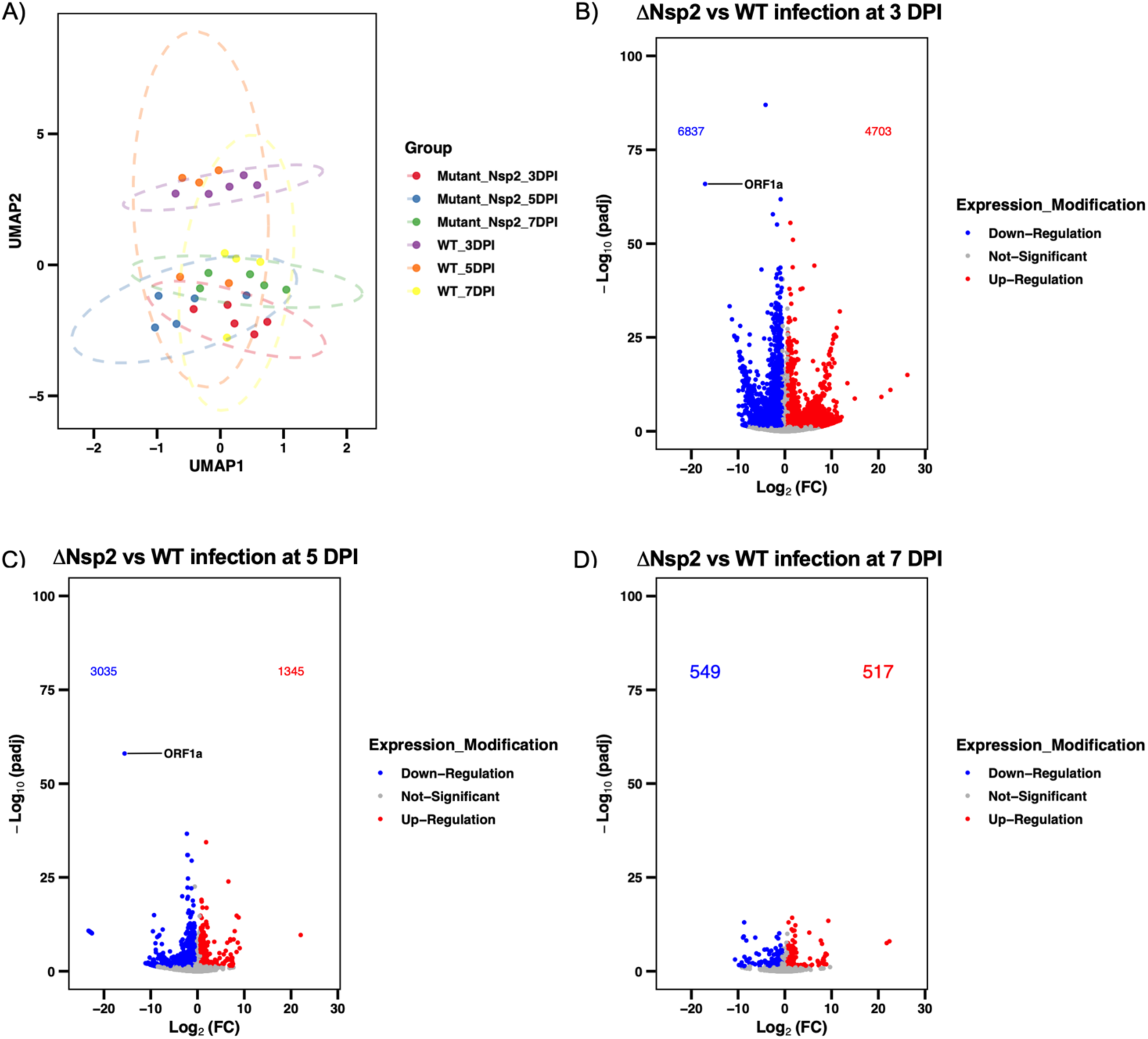
Differential regulation of the mice lung transcriptome following infection with wild-type or ΔNsp2 rSARS-CoV-2. (A) Dimension reduction of the transcriptome. Transcript counts were normalized using DESeq2 to stabilize variance, and Uniform Manifold Approximation and Projection (UMAP) was applied to visualize sample clustering and distances between conditions (B–D) Differential gene expression between ΔNsp2 and wild-type SARS-CoV-2-infected mice. Differential expression analysis was performed using DESeq2 from raw transcript counts. Results are presented as log2 fold change (log2FC) for ΔNsp2 relative to wild-type infection at 3 days post-infection (DPI) (B), 5 DPI (C), and 7 DPI (D) (mean; n = 4–5 per group). The y-axis represents the adjusted p-value (padj), corrected for multiple testing using a false discovery rate (FDR) threshold of 0.05. Transcripts with a padj below 0.05 and an absolute log_2_(FC) above 1.5 were colorized in red (up-regulated) or blue (down-regulated).

Across the infection course, comparison of ΔNsp2 and wild-type infections revealed a predominance of significantly downregulated transcripts in the ΔNsp2 condition (Figure 7B–D). This effect was most pronounced at early time points, consistent with the separation observed in the UMAP analysis. Notably, when each condition was compared to mock-infected controls, the proportion of up- and downregulated genes differed by less than 9% (Supplementary Figure 4A–B), suggesting that the observed differences between viruses primarily reflect reduced transcriptional induction rather than active repression in the absence of Nsp2.

In addition, lower expression of several viral transcripts was observed in ΔNsp2-infected mice across all time points. This effect was particularly evident at 3 DPI (9/12 detected viral genes) and 7 DPI (5/12), indicating altered viral transcriptional dynamics despite comparable overall replication. As a result of Nsp2 deletion, fewer transcripts falling in Orf1a gene were found in the ΔNsp2-infected mice.

### ΔNsp2 infection is associated with delayed and distinct transcriptional programs

To further characterize these differences, functional enrichment analysis was performed using Gene Set Enrichment Analysis (GSEA) on DESeq2-derived statistics comparing ΔNsp2 and wild-type infections at each time point. Enrichment analysis was conducted using the MSigDB M2 curated gene set collection, encompassing genetic and chemical perturbation signatures as well as canonical biological pathways. Only pathways significantly altered across all time points were retained, resulting in four major clusters based on shared enrichment patterns (Figure 8).

**Figure 8.**
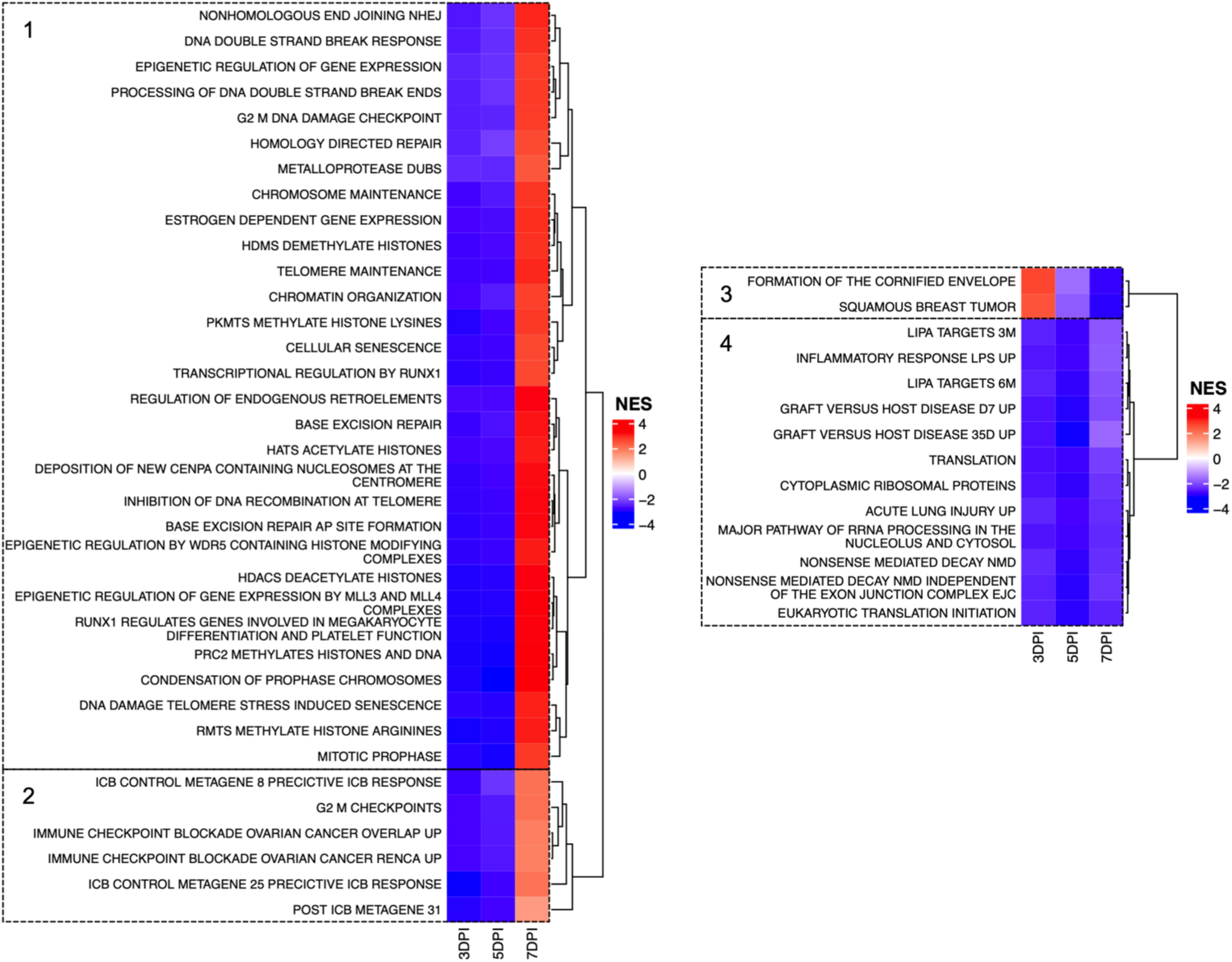
Functional enrichment analysis of lung transcriptomic changes in ΔNsp2 versus wild-type SARS-CoV-2 infection. Functional enrichment was performed using Gene Set Enrichment Analysis (GSEA) with the MSigDB M2 curated gene sets. DESeq2 Wald statistics from the ΔNsp2 versus wild-type comparison at each DPI were used as input for GSEA. The top 50 most differentially enriched pathways across all time points are shown in the heatmap. Pathways were hierarchically clustered based on enrichment patterns, and colors represent normalized enrichment scores (NES), with red indicating pathways enriched (upregulated) and blue indicating pathways depleted (downregulated) in ΔNsp2-infected mice relative to wild-type infection (n = 4–5 per group). The 4 major clusters were highlighted with dotted frame.

The first cluster included pathways related to DNA damage responses and epigenetic regulation, including histone modification. These pathways were strongly downregulated at 3–5 DPI, followed by a pronounced upregulation at 7 DPI, suggesting a delayed in the host response to the virus.

The second cluster comprised gene sets associated with successful immune checkpoint blockade [66, 67]. In ΔNsp2-infected mice, these pathways followed a similar trajectory but exhibited a more moderate upregulation at 7 DPI.

The third cluster displayed a distinct pattern, with early upregulation at 3 DPI followed by progressive downregulation at later time points. These gene sets were primarily associated with keratinocyte differentiation.

The fourth cluster encompassed pathways consistently downregulated throughout ΔNsp2 infection. These included gene sets related to RNA processing, translation, and acute lung injury/inflammatory responses. Notably, this cluster also contained gene signatures associated with lysosomal acid lipase (LIPA) deficiency [68].

### Nsp2 could modulate translation through RNA binding

To identify the underling mechanism behind Nsp2 dependant virulence, Cross-linking and Immunoprecipitation Sequencing (CLIP-seq) experiments were performed on A549-hACE2 cells infected with non-recombinant Wuhan-like SARS-CoV-2 (Figure 9A). Functional enrichment using ontology term were performed on the list of gene with peak identified by both peak callers (Figure 9B and Supplementary Figure 6). Genes identified were mainly related to the regulation of the translation machinery and RNA silencing process. Notably, those genes were also involved in the regulation of immunes response, metabolism and cell death. In support of the CLIP-seq results, proximity biotinylation experiments suggest that Nsp2 can interact with ribosomal proteins and member of the spliceosome complex (Supplementary Figure 7A and 7B).

**Figure 9.**
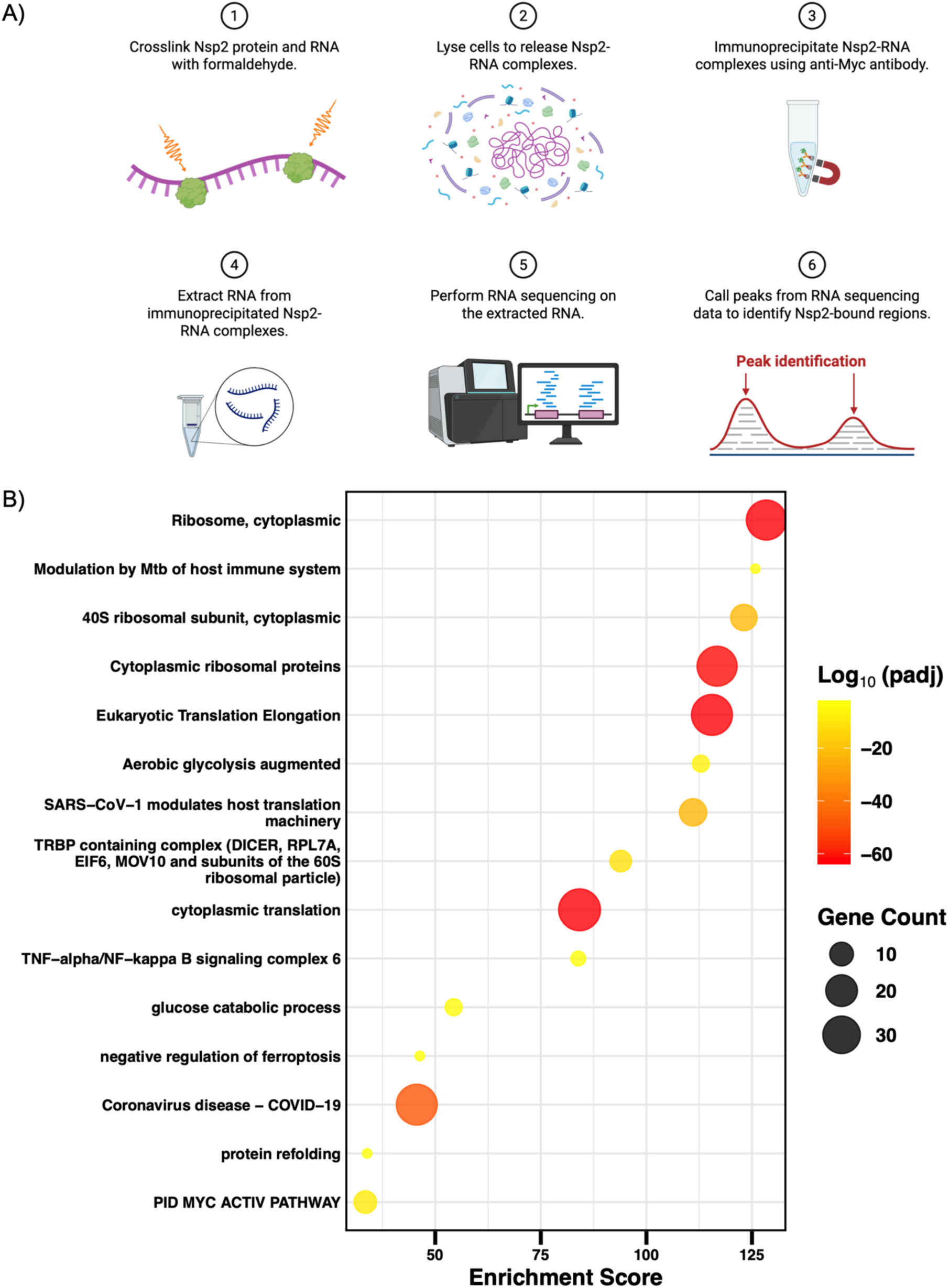
Identification of RNA interacting with SARS-CoV-2 Nsp2. (A) Schematic representation of Cross-linking and Immunoprecipitation Sequencing (CLIP-seq) procedure. The figure was created using BioRender. (B) Functional enrichment of transcript interacting with Nsp2. Genes with peak identified by both peak caller algorythms were analysed using Metascape. Term with a P adjusted value for false discovery rate (FDR) below 0.05 were retained. One representative term per cluster were presented and the top 15 are shown. Dot size represents the number of gene detected within the term (Gene Count), color scale represents the P adjusted value (padj) and x-axis represents the enrichment score.

## Discussion

As previously reported by our group using non-recombinant SARS-CoV-2, infection of K18-*hACE2* mice under our laboratory conditions was characterized by body weight loss, ruffled fur, reduced activity, and respiratory distress beginning approximately 5 days post-infection (DPI), with potential recovery by 10 DPI [37, 47, 51, 69, 70]. Other studies have reported fatal outcomes due to viral neuroinvasion following intranasal SARS-CoV-2 infection in K18-*hACE2* mice [71–73] However, under our experimental conditions, viral RNA in the brain was below the limit of detection or detected at very low levels in only a few mice ruling out encephalitis as a cause of death (Supplementary Figure 2).

Disease symptoms alike those observed with non-recombinant WT SARS-CoV-2 were phenocopied in mice infected with WT rSARS-CoV-2. However, mice infected with the ΔNsp2 mutant showed minimal weight loss and overall more favorable outcomes. This difference could not be explained by replication defects as ΔNsp2 mutant and wild type viruses replicated similarly both in vitro and in vivo. Although the WVPRA motif in Nsp2 sequence was reported to be required for efficient Nsp1 and Nsp3 self-cleavage [74], in the context of infection we did not observe any impairment of this process in the absence of Nsp2 under our experimental conditions.

A trend toward enhanced viral clearance was observed at 7 DPI in mice infected with the ΔNsp2 virus, suggesting a more effective antiviral response. Xu and colleagues reported that Nsp2 inhibits IFNβ protein production [25]; we did not observe increased type I interferon production or in an increase in ISGs expression in the lungs of ΔNsp2-infected mice in comparison with their WT-infected counterparts.

Compared to wild-type virus, infection with ΔNsp2 mutant virus was associated with reduced lung and systemic inflammation. Higher levels of IL-6 and IFNγ were detected in the lungs of mice infected with wild-type rSARS-CoV-2, consistent with the more severe disease observed in this group. These findings align with reports showing elevated inflammatory cytokines in bronchoalveolar lavage fluid (BALF) from patients with poor COVID-19 outcomes [75, 76]. Moreover, gene expression signature associated with acute inflammation and acute lung injury were reduced in ΔNsp2-infected mice, further supporting these observations. Similarly, increased levels of IFNγ, CCL2, and CXCL10 (IP-10) in the blood have been associated with severe COVID-19 in humans [77–79], consistent with our observations in mice infected with the wild-type virus. In addition, gene signatures associated with lysosomal acid lipase (LIPA) knock-down were downregulated in the lungs of ΔNsp2-infected mice, which may contribute to the modulation of inflammatory responses [80, 81]. These findings are consistent with the established role of bioactive lipids in SARS-CoV-2 infection and inflammation [18, 19, 37, 47, 51].

Transcriptional signatures related to DNA damage responses were suppressed at early time points in ΔNsp2-infected mice. SARS-CoV-2 infection has previously been associated with the accumulation of cytoplasmic chromatin and increased levels of circulating cell-free DNA (cfDNA) [82, 83]. This accumulation of damaged DNA can activate inflammatory pathways through pattern recognition receptors (PRRs) [7, 82, 84]. Therefore, reduced activation of DNA damage–associated pathways in ΔNsp2 infection may contribute to the attenuated inflammatory response observed in this model.

Notably, although most inflammatory mediators were elevated in wild-type–infected mice, CCL5 production was more pronounced in the lungs of ΔNsp2-infected mice. Increased CCL5 levels in the lungs have been associated with improved outcomes following influenza infection, promoting lymphocyte recruitment and providing anti-apoptotic signals to macrophages [85, 86]. These functions are consistent with the increased abundance of macrophages and lymphocytes observed in the lungs at 7 DPI in ΔNsp2-infected mice. In patients with COVID-19, lower plasma levels of CCL5 have been associated with more severe disease [87, 88]. In our study, circulating CCL5 levels were close to the limit of detection; however, a trend toward increased levels was observed at 7 DPI in wild-type–infected mice. Additionally, reduced CCL5 expression in the upper respiratory tract has been linked to increased disease severity in COVID-19 patients, which is consistent with the higher pulmonary production of CCL5 observed in ΔNsp2-infected mice [89].

Severe COVID-19 has been strongly associated with peripheral lymphopenia [90, 91]. Accordingly, compared with ΔNsp2 infection, lower proportions of T, B, and NK cells were detected in both the blood and lungs of mice infected with wild-type rSARS-CoV-2 at 5 and 7 DPI. This reduction, together with the increased abundance of monocytes in wild-type–infected mice, suggests that Nsp2 may skew the immune response toward a pathogenic inflammatory profile. An early Th1 response was observed in the lungs of wild-type–infected mice. While Th1 responses have been associated with favorable COVID-19 outcomes [92, 93], excessive type I immune responses may also contribute to hyperinflammation [22, 94].

Interestingly, transcriptomic signatures associated with immune checkpoint blockade were suppressed at early time points in ΔNsp2-infected mice but became upregulated at 7 DPI. This pattern suggests a delayed engagement of immune regulatory mechanisms, potentially allowing for a more controlled early antiviral response followed by effective resolution at later stages. This temporal shift may be linked to the concurrent modulation of pathways involved in gene expression regulation, which exhibited a similar dynamic profile.

Infection with the ΔNsp2 mutant was further characterized by a marked shift in antigen presentation dynamics. Conventional and plasmacytoid dendritic cells (cDC and pDC) were less abundant, whereas interstitial macrophages (IMΦ), megakaryocytes (MK), and CD11c⁺ BST2⁺ Ly6C⁺ monocytes expressing MHC class II were present at higher proportions in the lung. Taken together, these results suggest that a broader array of immune cells may efficiently present antigens and promote a more effective T-cell response. Given that SARS-CoV-2 infection of monocytes, macrophages, T cells, and NK cells has been reported [95–97], it is plausible that Nsp2 directly modulates the function and phenotype of these leukocyte populations.

A reduction in total leukocyte numbers, along with decreased proportions of Th, CTL, and B cells in the blood at 7 DPI in wild-type–infected mice, further supports the presence of virus-induced lymphopenia, a feature not observed following ΔNsp2 infection. The association between lymphopenia and COVID-19 severity has been extensively documented [98–100]. Proposed mechanisms include exposure to cytokine storm mediators (IL-1β, IL-6, TNFα, and IFNγ) and interactions with myeloid-derived suppressor cells (MDSCs) [101, 102]. Although elevated IFNγ levels were detected in the plasma of wild-type–infected mice, other cytokines were below the limit of detection, potentially due to BSL-3 sample inactivation procedures. In mice, polymorphonuclear-MDSCs and monocytic-MDSCs (M-MDSCs) are defined as CD11b⁺Ly6G⁺Ly6C^lo^ and CD11b⁺Ly6G⁻Ly6C^hi^ cells, respectively, though distinguishing these populations from other myeloid cells remains challenging [102]. In humans, HLA-DR expression can be used to discriminate monocytes from M-MDSCs [102]. We therefore hypothesize that the altered MHC class II expression observed in CD11c⁺ BST2⁺ Ly6C⁺ monocytes in the lungs of wild-type–infected mice may reflect an increase in M-MDSCs.

A trend toward peripheral neutrophilia was observed in response to wild-type infection but not following ΔNsp2 infection. Neutrophilia has been associated with COVID-19 severity and mortality [100]. Interestingly, higher proportions of PD-L1–expressing neutrophils were detected in the blood of ΔNsp2-infected mice at 3 DPI. As PD-L1 is known to exert immunosuppressive effects [103], these neutrophils may contribute to limiting excessive inflammation and exert a protective role, as described in tuberculosis [104].

As a potential mechanism of action, SARS-CoV-2 Nsp2 has been reported to modulate host translation [25, 26, 105]. In line with this, we observed reduced expression of gene sets associated with translation and RNA processing in ΔNsp2-infected mice. Given that Nsp2 contains a predicted zinc-finger motif, it is plausible that this protein may directly bind RNA and influence these pathways [106–108]. Supporting this hypothesis, transcripts from several gene have been identified by CLIP-seq on Nsp2 in the context of infection. The role of those genes identified combined with mouse RNA-seq data suggest that Nsp2 may interact with host translational and post-transcriptional machinery to modulate cellular functions in a manner that promotes its virulence.

This study has several limitations. First, we did not identify the precise molecular mechanism underlying Nsp2-dependent SARS-CoV-2 pathogenesis. However, our results provide strong in vivo evidence of its role and generate testable hypotheses for future mechanistic studies. Second, due to biosafety level 3 (BSL-3) constraints and associated costs, the number of animals per group was limited, which may reduce statistical power, the generalizability of the findings and the detection of sex differences. Finally, although the K18-hACE2 mouse model recapitulates key features of severe COVID-19, differences between murine and human immune responses, along with the non-physiological expression pattern of hACE2, may limit direct extrapolation of these results to human disease.

In summary, this study demonstrates that SARS-CoV-2 Nsp2 contributes to viral pathogenesis in vivo. Deletion of Nsp2 resulted in a marked attenuation of disease severity, associated with reduced pulmonary and systemic inflammation. This attenuation was accompanied by a shift in leukocyte dynamics, from a pro-inflammatory profile in wild-type infection toward a more balanced and effective antiviral response in ΔNsp2-infected mice. Lung transcriptomic, Chip-seq and BioID analyses further suggest that Nsp2 influences host responses through modulation of RNA processing and translation-related pathways. Collectively, these findings identify Nsp2 as a key regulator of SARS-CoV-2–induced immunopathology and provide new insights into host–pathogen interactions in a physiologically relevant in vivo model.

## Acknowledgments

We thank the high throughput image analysis and flow cytometry core facilities of Centre de Recherche du CHU de Quebec for providing access to the slide scanner and cytometer used in this study. We also thank the Sanger and NGS core facilities of Centre de Recherche du CHU de Quebec for sequencing services. We thank the Plateforme Protéomique du Centre de Génomique de Québec for their mass-spectrometry protein identification service. We also thank Dr Vladimir Larionov for providing the VL6-48N yeast strain. Graphical abstract, Figure 1A, Figure 2A and Figure 8A were produced using Biorender. ChatGPT 5.2 (OpenAI) was used for improving the text readability. Finally, we thank Julien Prunier for providing bioinformatics training.

## Founding

This research was enabled by support provided by a CIHR operating grant to the Coronavirus Variants Rapid Response Network (CoVaRR-Net) (ARR-175622) and a CIHR Operating Grant: Emerging COVID-19 Research Gaps and Priorities Funding Opportunity awarded to LF.

## Author contributions

Study conceptualization, methodology, resources, project administration and supervision was performed by EL, LF, ID and AG. EL and LF were responsible to writ the original draft. EL, ID, JL, AG, LG, CJB, MF, PF and MRB contribute to investigation, validation and data curation. EL, ID, CJB, MRB and LF performed formal analysis. Funding acquisition was performed by LF and AD. All the authors contribute to review and editing of the manuscript.

## Declaration of generative AI and AI-assisted technologies in the manuscript preparation process

During the preparation of this work the authors used ChatGPT 5.2 (OpenAI) to improving the text readability and grammar and spelling check. BioRender AI tool to inspire the graphical abstract design. After using this tool/service, the authors reviewed and edited the content as needed and takes full responsibility for the content of the published article.

## Institutional review board

The study was conducted in accordance with the Declaration of Helsinki, and Mice protocols were approved by the Comité de protection des animaux de l’Université Laval. (Approval code: 2023-1292).

## Materials availability

Plasmids and cell lines are available from the lead contact with a completed materials transfer agreement.

## Data and code availability

Raw data and code generated in this study will be shared upon request the lead contact after publication.

