## Supplementary tables and figures for "SARS-CoV-2 Nsp2 reprograms host immunity to drive pathogenic inflammation"

**Supplementary Table 1.** Antibody and streptavidin used for western blot assay.

| Antibody | Working Concentration | Supplier/cat | Clone |
|---|---|---|---|
| Rabbit anti–SARS-CoV-2 Nsp1 | 0.6 µg/mL | Abclonal (A20200) | Polyclonal |
| Rabbit anti–SARS-CoV-2 Nsp2 | 0.25 µg/mL | Abcam (ab284036) | EPR24853-4 |
| Rabbit anti–SARS-CoV-2 Nsp3 | 52 ng/mL | Cell Signaling Technology (88086) | Polyclonal |
| Rabbit anti–SARS-CoV-2 N | 0.2 µg/mL | Rockland Immunochemicals Inc. (200-401-MS4) | Polyclonal |
| Mouse anti–SARS-CoV-2 S | 1 µg/mL | Abcam (ab272504) | Polyclonal |
| Mouse anti-GFP Tag | 1 µg/mL | Applied Biological Materials (G096) | LGB-1 |
| Mouse anti-HA Tag | 1.11 µg/mL | - | 12CA5 |
| Alexa Fluor™ 488–conjugated goat anti-rabbit IgG | 0.5 µg/mL | Thermo Fisher (A11034) | Polyclonal |
| Alexa Fluor™ 647–conjugated Donkey anti-rabbit IgG | 0.5 µg/mL | Jackson ImmunoResearch Laboratories Inc. (711-606-146) | Polyclonal |
| Alexa Fluor™ 488–conjugated goat anti-mouse IgG | 0.5 µg/mL | Jackson ImmunoResearch Laboratories Inc. (115-546-003) | Polyclonal |
| Alexa Fluor™ 647–conjugated goat anti-mouse IgG | 0.5 µg/mL | Jackson ImmunoResearch Laboratories Inc. (115-606-146) | Polyclonal |
| Peroxidase Donkey Anti-Mouse IgG | 40 ng/mL | Jackson ImmunoResearch Laboratories Inc. (715-035-151) | Polyclonal |
| Peroxidase Streptavidin | 50 ng/µL | Jackson ImmunoResearch Laboratories Inc. (016-030-084) | - |

**Supplementary Table 2.** Leucocyte staining antibody cocktail.

| Marker | Clone | Fluorochrome | Supplier | Dilution Lung | Dilution Blood |
|---|---|---|---|---|---|
| CD16/CD32 (Mouse Fc block) | 2.4G2 | N.A. | BD Biosciences | 1μg | 2μg |
| CD3 | 17A2 | Brilliant Violet 421 | Thermo Fisher | 800ng | 500ng |
| CD161/NK1.1 | PK136 | Brilliant Violet 480 | Thermo Fisher | 500ng | 500ng |
| CD317/BST2 | 927 | Brilliant Violet 605 | Biolegend | 500ng | 500ng |
| MHCII(I-A/I-E) | M5/114.15.2 | Brilliant Violet 650 | Thermo Fisher | 200ng | 200ng |
| CD119/IFNGR1 | GR20 | Brilliant Violet 711 | BD Biosciences | 500ng | 500ng |
| F4/80 | BM8 | Brilliant Violet 785 | Biolegend | 1μg | - |
| CD8α | 53-6.7 | FITC | BD Biosciences | 750ng | 750ng |
| CD41 | MWReg30 | RealBlue 545 | BD Biosciences | 250ng | 250ng |
| CD4 | RM4-5/ RM4.5 | PE | BD Biosciences | 150ng | 150ng |
| Ly-6C | AL-21 | PE-CF594 | BD Biosciences | 50ng | 50ng |
| CD11c | HL3 | RealBlue 705 | BD Biosciences | 200ng | 200ng |
| CD274/PD-L1 | MIH5 | RealBlue 780 | BD Biosciences | 300ng | 300ng |
| Ly-6G | 1A8 | APC | BD Biosciences | 400ng | 400ng |
| CD45 | 30-F11 | APC-Cy7 | BD Biosciences | 50ng | 50ng |
| CD45R/B220 | RA3-6B2 | eFluor 506 | Thermo Fisher | 1μg | 500ng |
| CD11b | M1/70 | Alexa Fluor 660 | Thermo Fisher | 250ng | 250ng |

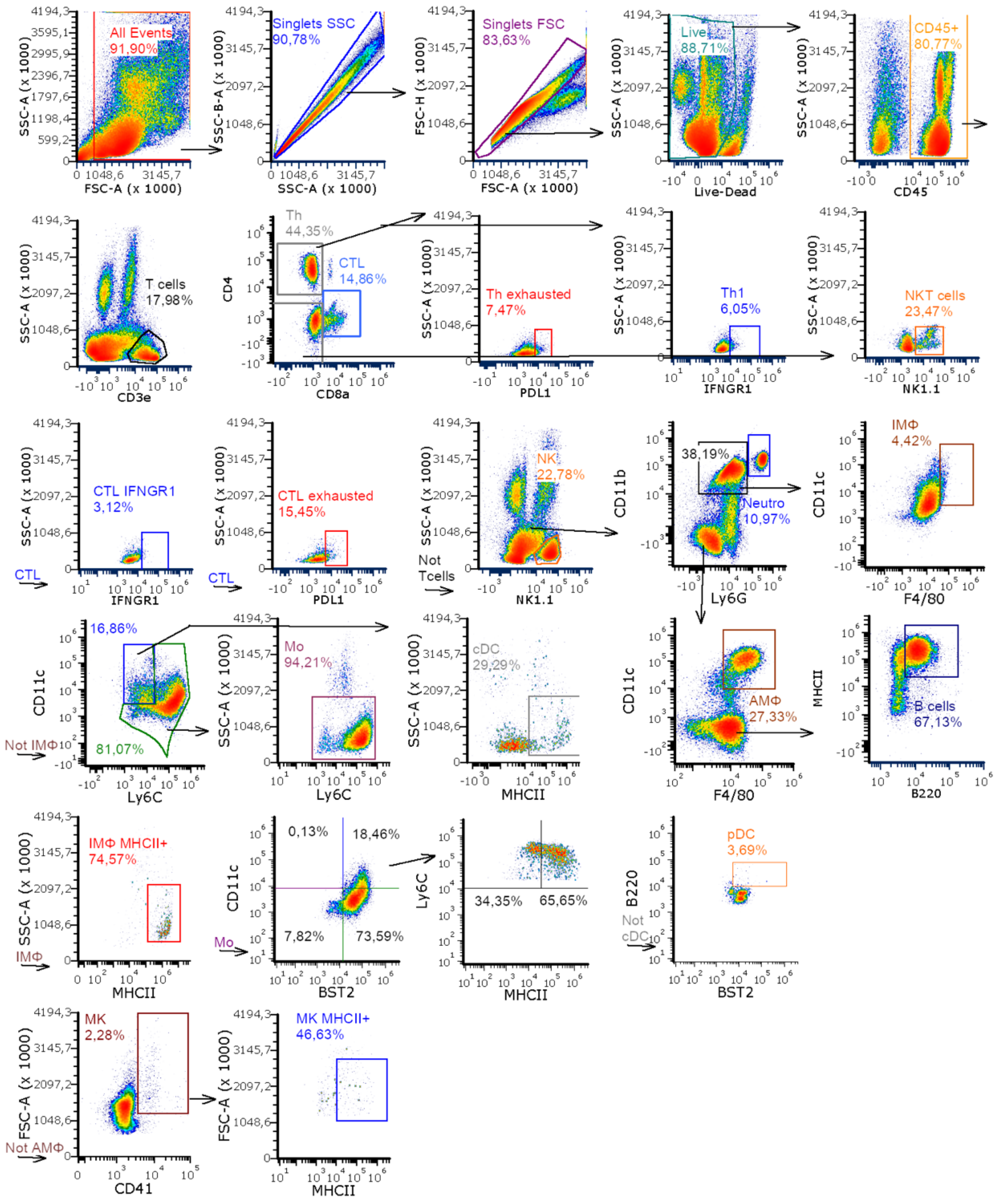


**Supplementary figure 1A. Gating strategy for lung immunophenotyping.** Density plots illustrate the gating strategy used to identify each leukocyte subpopulation in lung tissue. For most populations, a representative mouse at 3 days post-infection with the ΔNsp2 mutant virus is shown. For alveolar macrophages (AMΦ) and megakaryocytes (MK), a representative mouse from the mock-infected group was used. Population abbreviations are as follows: T helper (Th) cells, cytotoxic T lymphocytes (CTL), T helper type 1 (Th1) cells, interstitial macrophages (IMΦ), natural killer (NK) cells, monocytes (Mo), neutrophils (Neutro), classical dendritic cells (cDC), and plasmacytoid dendritic cells (pDC).

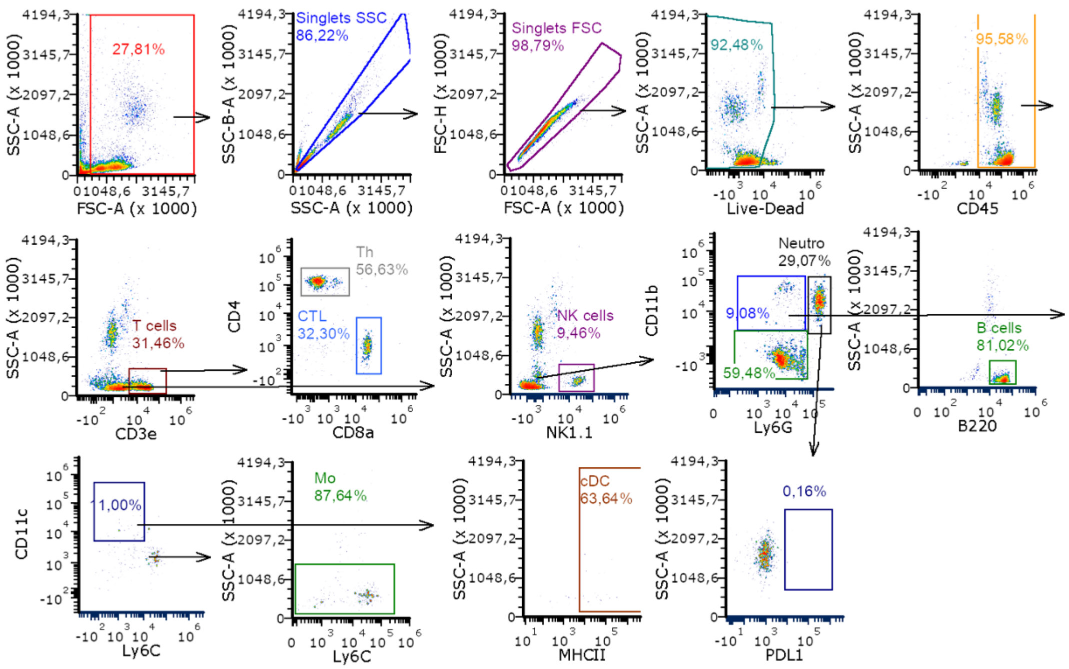


**Supplementary figure 1B. Gating strategy for blood immunophenotyping.** Density plots indicate the gating strategy to identify each leucocyte sub-population. A representative mouse from mock group was used for each graph.

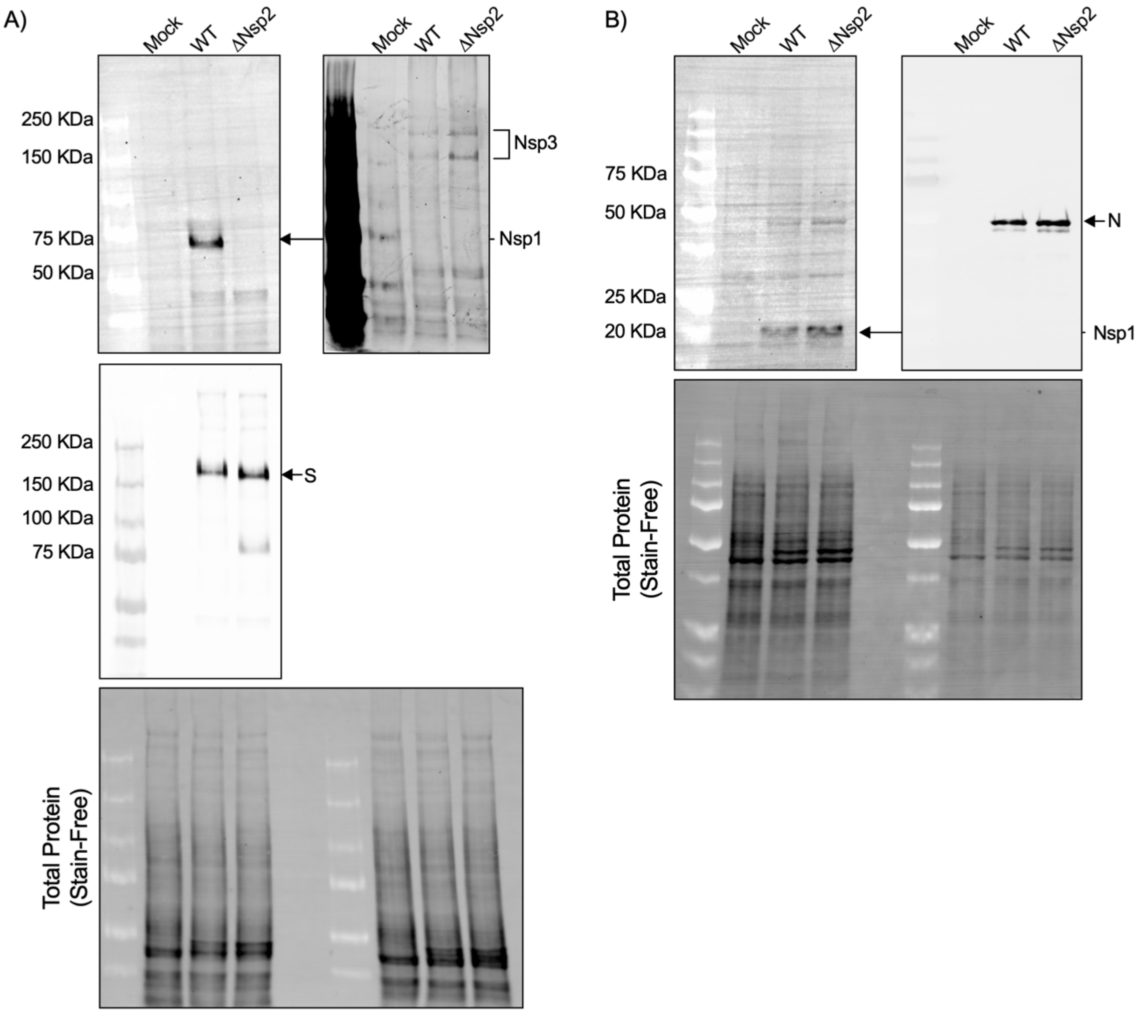


**Supplementary figure 4. Uncropped Western blot and loading control for viral protein expression in Vero cells.**
Uncropped images of membranes probed with antibodies against SARS-CoV-2 Nsp2, Nsp3, and S (A), or SARS-CoV-2 Nsp1 and N (B). Total protein staining using Stain-Free technology is shown in the lower part of each panel as a loading control.

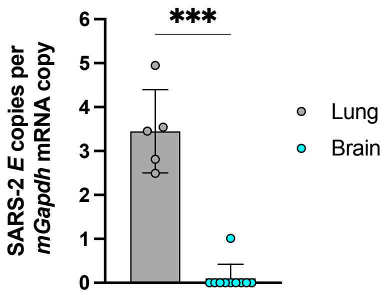


**Supplementary figure 3. Comparison of the viral RNA detection in the lung and brain of K18-*hACE2* mice.**

Total RNA was extracted from lung and brain tissues of mice infected with wild-type rSARS-CoV-2 on 5 DPI, as described in the Methods section. SARS-CoV-2 *E* RNA was quantified by RT-ddPCR and expressed as copies of viral *E* RNA per copy of mouse *Gapdh* mRNA. Data are presented as mean ± SD (n = 5 for lung and n = 10 for brain). Viral RNA levels between organs were compared using a two-tailed Mann–Whitney test. ***P < 0.0002.

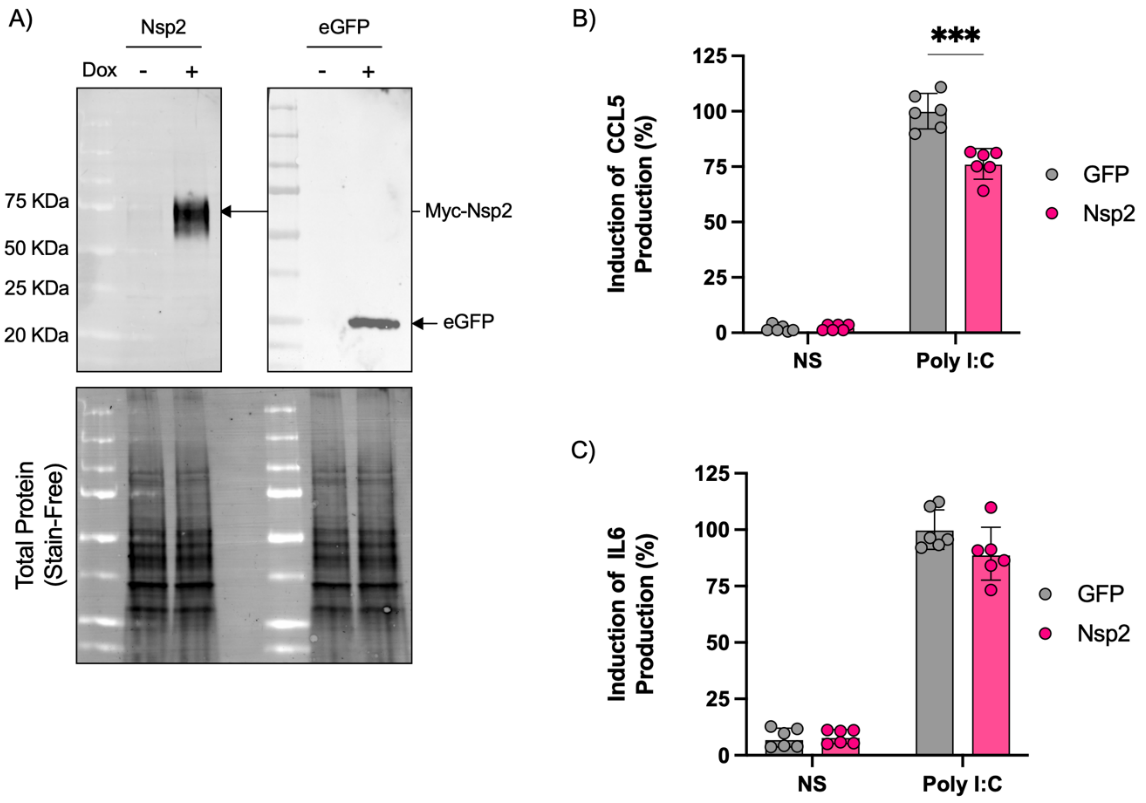


**Supplementary figure 4. In vitro inhibition of the CCL5 production by SARS-CoV-2 Nsp2.**
(A) Induction of transgene expression in A549-hACE2 GFP and SARS-CoV-2 Nsp2-expressing cell lines. Cells were induced and harvested as described in the Materials and Methods section. Protein expression was analyzed by Western blot using antibodies against SARS-CoV-2 Nsp2 (AF488) or GFP tag (AF488). Stain-Free technology was used as a loading control.
(B–C) Induction of CCL5 (B) and IL-6 (C) production in cell supernatants following transgene induction and Poly I:C transfection in A549-hACE2 GFP and SARS-CoV-2 Nsp2-expressing cell lines. Supernatants were collected and cytokine levels were quantified by ELISA as described in the Materials and Methods section. Data are presented as relative induction, normalized to the corresponding GFP control (set to 100%) (mean ± SD; n = 6 per group). Statistical comparisons between GFP and Nsp2-expressing cells were performed using a log-normal t-test. ***P < 0.0002.

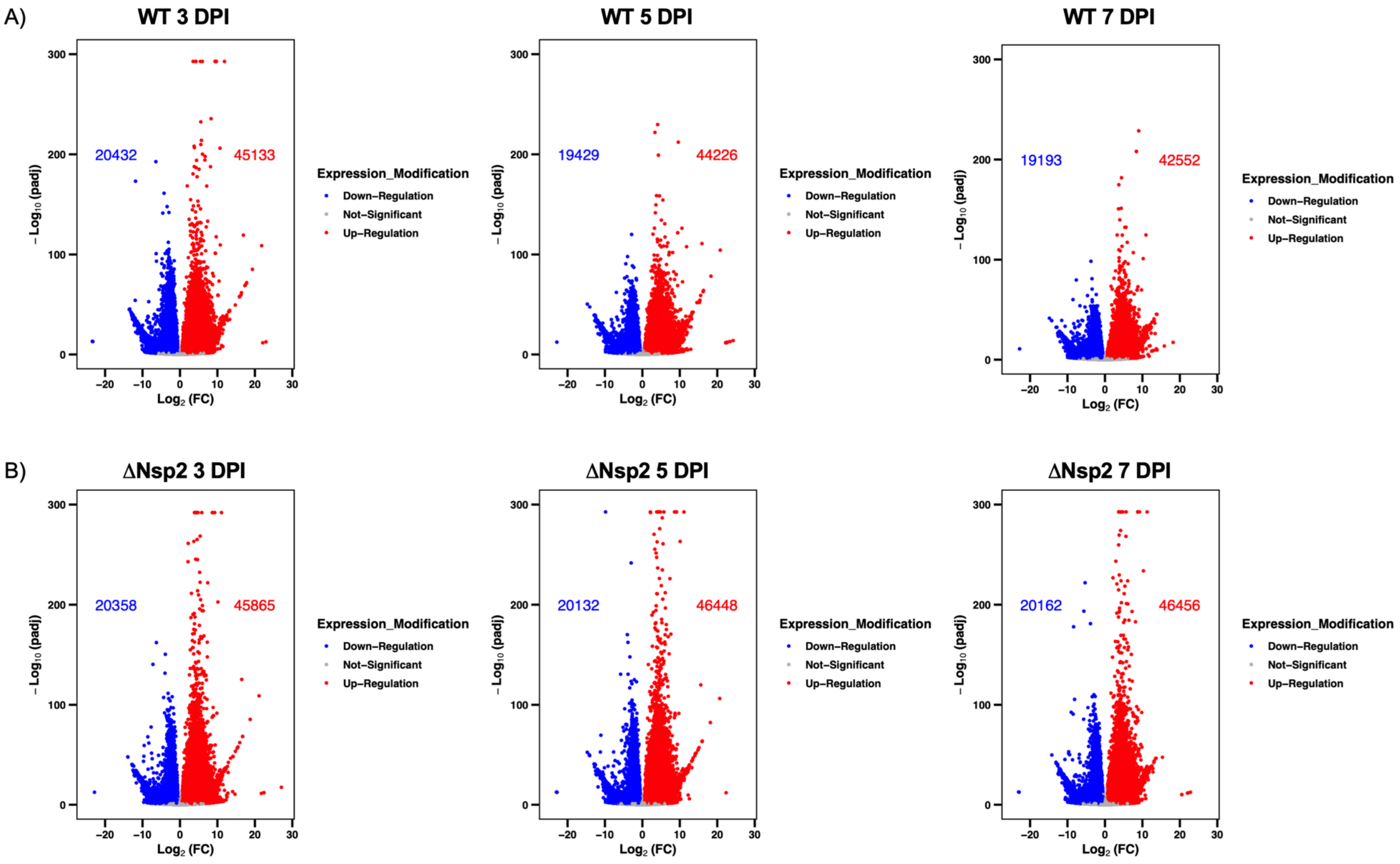
A)
WT 3 DPI
20432
45133
WT 5 DPI
19429
44226
WT 7 DPI
19193
42552
B)
ΔNsp2 3 DPI
20358
45865
ΔNsp2 5 DPI
20132
46448
ΔNsp2 7 DPI
20162
46456
−Log10 (padj)
Log2 (FC)
Expression_Modification
Down−Regulation
Not−Significant
Up−Regulation

**Supplementary figure 5. Lung transcriptional change following infection with wild-type or ΔNsp2 rSARS-CoV-2.**

From transcript raw quantification, statistical change between infected mice were evaluated using DESeq2. Data were shown in $\log_2$ fold of change ($\log_2$[FC]) between rSARS-CoV-2 wild-type (A) or ΔNsp2 (B) and mock infected mice. For each transcript, P value adjusted (padj) to a false discovery rate (FDR) of 0.05 value was display in Y axis and transcripts with a padj below 0.05 and an absolute $\log_2$(FC) above 1.5 were colorized in red (up-regulated) or blue (down-regulated).

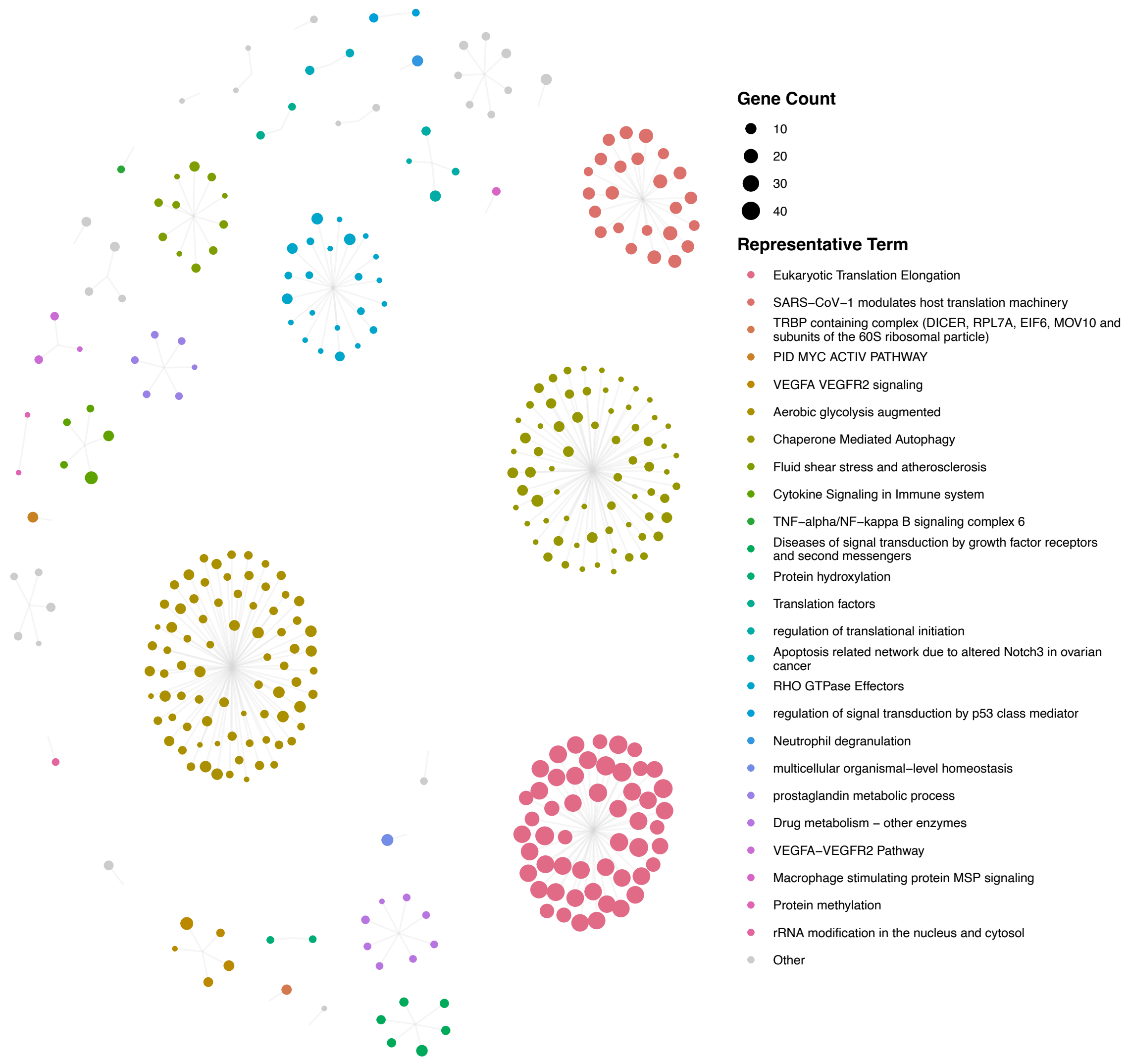


**Supplementary figure 6. Functional enrichments of gene with Nsp2 CLIP-seq peak.**

Genes with peak identified by both peak callers were analysed using Metascape. Term with a P adjusted value (FDR) below 0.05 were retained. One representative term per cluster were presented. Dot size represents the number of gene detected within the term (Gene Count) and color were associated with each representative term.

A)

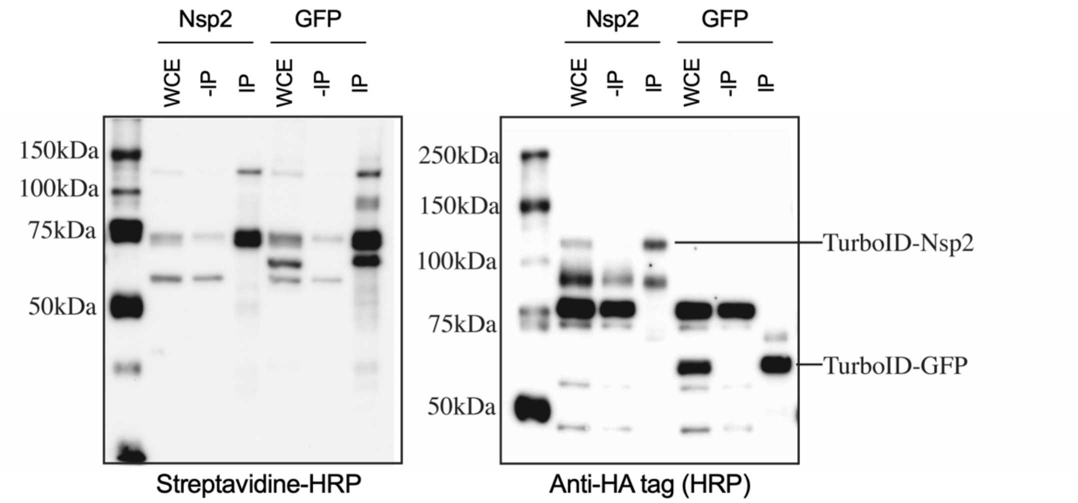


B)

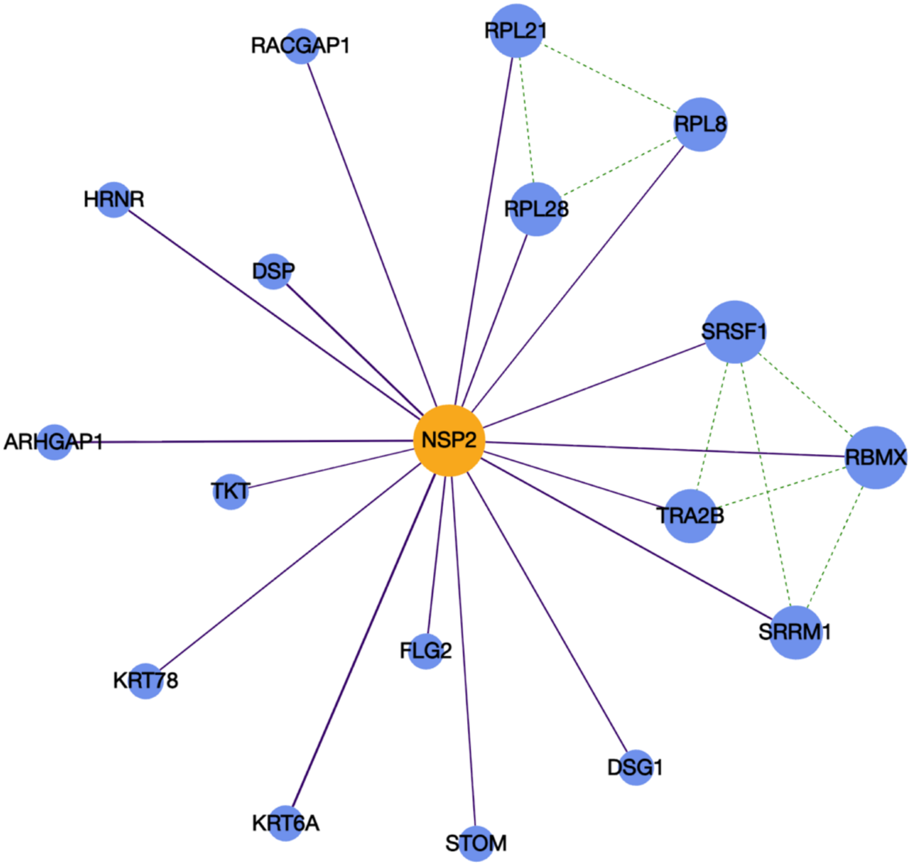


**Supplementary figure 7. Proximity biotinylation of presumptive Nsp2 interactors.**

(A) Expression of fusion proteins and enrichment of biotinylated protein. Efficiency of streptavidin pull-down and fusion protein expression were evaluated by western bolt using HRP-labeled Streptavidin (Left) or antibodies against HA tag (HRP) (Right).

(B) Network of protein-protein interaction identified with Nsp2 proximity biotinylation. Peptide count obtained from mass-spectrometry analysis of pull-down protein were analysed with SAINT using BioGRID database to map know interactions. Protein with a SAINT score below 0.6 and a fold change A (FCA) above 1 were shown. Purple line indicate interaction with Nsp2 when green dashed line indicate interaction reported in BioGRID.
